# mRNA-Encoded TLR5 Agonist as an Immune Adjuvant

**DOI:** 10.64898/2026.05.30.728968

**Authors:** Alexandra V. Vylegzhanina, Oscar Murillo Gomez, Ilya I. Gitlin, Ivan Molodtsov, Anatoli S. Gleiberman, Kamila Agadilova, Nayeli Molina Acevedo, Ekaterina L. Andrianova, Benjamin Israelow, Andrei V. Gudkov

**Affiliations:** Flag, Bio, Inc., Buffalo, NY, USA; Yale University School of Medicine, New Haven, CT, USA; Roswell Park Comprehensive Cancer Center, Buffalo, NY, USA

## Abstract

Activation of innate immunity enhances adaptive immune responses and underlies the mechanism of action of vaccine adjuvants. First-generation mRNA vaccines lack a dedicated adjuvant, relying instead on the self-adjuvanting properties of lipid nanoparticles (LNPs), and are thus expected to benefit from additional engagement of innate immune pathways not activated by LNPs. Among innate immune modulators, TLR5 agonists are particularly promising adjuvants due to their ability to induce a balanced Th1/Th2 immune response and their relatively favorable safety profile. Here we tested whether supplementing mRNA vaccines with mRNA encoding a TLR5 agonist could enhance immunization efficacy by induction of TLR5 signaling coordinated with vaccine antigen expression. We designed FL711, an mRNA encoding a derivative of Salmonella flagellin optimized for mammalian expression, functionally active in TLR5 signaling, deimmunized for pre-existing human T and B cell epitopes, and engineered for secretion to stimulate the TLR5 pathway in the local tissue microenvironment. We characterized FL711 in vitro and in vivo for functional and pharmacological parameters and assessed its adjuvant effect as a component of experimental anti-influenza and anti-SARS-CoV-2 mRNA vaccines. Supplementation with small amounts of FL711 mRNA (up to 30-fold less than antigen-encoding mRNA) significantly enhanced vaccine immunogenicity and protective efficacy, stimulating local NF-κB induction, boosting antibody production and T cell activation, and prolonging the durability of the response — while enabling a marked reduction in mRNA dose per vaccine. These findings support the potential of FL711 as a broadly applicable mRNA-encoded adjuvant to improve the potency, durability, and dose efficiency of next-generation mRNA vaccines.

## Introduction

Vaccine adjuvants play a central role in shaping the magnitude, quality, and durability of immune responses. Their effects extend beyond simple enhancement of immunogenicity and include prolongation of antigen availability, improved delivery of antigen to draining lymph nodes, activation and maturation of antigen-presenting cells, and programming of the cytokine milieu required for effective adaptive immunity. Mechanistically, adjuvants can act either as direct ligands of pattern-recognition receptors (PRRs) or as inducers of cellular stress and death that trigger the release of damage-associated molecular patterns and other endogenous immunostimulatory signals. These functions have become increasingly important as vaccination has expanded beyond classical prophylaxis against infectious diseases toward more demanding applications, including vaccines for cancer and other conditions in which distinct types of immune polarization, durability, and tissue targeting may be required (review (*1*)).

Modern adjuvant development was transformed by the discovery of pathogen-associated molecular patterns and the elucidation of innate immune pathways linking PRR activation to the induction of adaptive immunity, particularly through dendritic cell activation (*2*). Yet despite the large number of experimental PRR ligands described over the past two decades, only a limited number have been incorporated into licensed vaccines, most often in combination with older delivery systems such as alum (*3*) or liposomal formulations (*4*). A prominent example is the AS04 adjuvant system, which combines alum with 3-O-desacyl-4′-monophosphoryl lipid A (MPLA), a detoxified TLR4 agonist derived from lipopolysaccharide. AS04 is used in the licensed vaccines Cervarix and Fendrix, and clinical experience with these vaccines has shown that AS04 can improve the magnitude and durability of antibody responses relative to alum-only comparators in settings where stronger immunogenicity is needed, including populations at elevated risk of hepatitis B virus infection and in human papillomavirus vaccination (*5*). Furthermore, the example of AS04, highlights the increasing recognition that combination adjuvants acting on multiple PRRs can enhance vaccines responses though synergistic effects. Exploring TRL agonists as prospective vaccine adjuvants continues with TLR9 agonist CpG 1018 recently approved for the hepatitis B vaccine (*6*), (*7*), while TLR7/8 agonists continue to attract major interest as next-generation immunostimulatory components (*8*), (*9*), (*10*).

There is a clear unmet need for improved vaccines against respiratory viruses, given the continuing morbidity, mortality, healthcare, and economic burden due to epidemic or endemic respiratory viruses. The rapid deployment of LNP(mRNA) vaccines during the COVID-19 pandemic underscores the strategic value of platform technologies that are antigen-agnostic and enable rapid manufacturing scale-up, thereby accelerating responses to emerging pathogens (*11*). However, the transition to mRNA vaccines has introduced a new challenge for adjuvant design: innate immune activation must be triggered in the same site and within the same time window as antigen expression. In current LNP(mRNA) vaccines, this requirement is met only partially by the immunostimulatory properties of the lipid nanoparticle carrier (*12*). This solution is clearly effective enough to support highly protective vaccines, but it is not necessarily optimal (*13*). Recent mechanistic work further strengthens this view by showing that some ionizable lipids may be able to signal through TLR4 to activate NF-κB and IRF pathways (*14*). Innate sensing of the mRNA component is a double-edged sword, because excessive activation can suppress antigen expression through interferon-linked pathways involving PKR and OAS, thereby shifting the balance from productive antigen synthesis toward inflammation. At the same time, the LNP component, particularly the ionizable lipid, acts as the dominant source of adjuvanticity by inducing inflammatory cytokines and chemokines, promoting antigen-presenting-cell activation, and supporting IL-6-dependent T follicular helper and germinal-center responses as well as type I interferon-associated cytotoxic T cell responses (*12*). However, these signals are broad, only partially controllable, and not necessarily aligned with the immune phenotype required for a given application; they also contribute to vaccine reactogenicity, including fever and local inflammatory reactions (*15*).

Consistent with this, rational redesign of LNPs with TLR7/8-agonistic adjuvant lipidoids (*16*), multiply adjuvanted LNP formulations (*17*), IL-12 co-administration with LNP(mRNA) (*18*), and mRNA-encoded constitutively active STING(V155M) (*19*) have all been shown to enhance specific immune responses beyond those achieved by conventional LNP(mRNA) formulations. Together, these findings suggest that the next generation of mRNA vaccines will require finer control over innate immune activation to preserve efficient antigen expression, minimize unnecessary reactogenicity, and direct the magnitude, durability, and polarization of adaptive immunity appropriate for each vaccine setting.

Among innate immune receptor agonists, flagellin is uniquely suited for incorporation into gene-based vaccines because, unlike most such agonists, it is itself a protein ligand of TLR5 and can therefore be genetically encoded (*20*). This feature enables adjuvant and antigen to be expressed in the same cells and within the same temporal window, an especially attractive property for mRNA vaccine platforms. Experimental studies in DNA- and mRNA-based systems support this strategy, showing that encoded flagellin can enhance vaccine immunogenicity, particularly cellular immune responses (*21*), (*22*), (*23*). Flagellin has also repeatedly shown potent adjuvant activity, often inducing a mixed or balanced Th1/Th2 response profile rather than the more polarized Th1 response that current LNP(mRNA) vaccines produce (*24*). Notably, the TLR5 pathway appears relatively resistant to age-associated decline, and flagellin-based influenza vaccine candidates have demonstrated favorable tolerability and strong serologic immunogenicity in mice and older adults (*25*). Collectively, these observations provide a strong mechanistic and translational rationale for using TLR5 agonists, including mRNA-encoded flagellin, as vaccine adjuvants. In our prior work, we developed a deimmunized TLR5 agonist optimized for human use that avoids preexisting neutralizing antibody and T-cell responses (*26*), providing an attractive candidate for mRNA-encoded delivery. In the present study, we designed and generated a synthetic mRNA, named FL711, encoding a structurally optimized and deimmunized flagellin derivative with enhanced expression, functionality, and secretion in mammalian systems. We then evaluated FL711 in vitro and in vivo as an immune adjuvant in the context of two experimental antiviral mRNA vaccines and found that it substantially improved antibody responses, T cell activation, durability, and protective efficacy, and overall immunization efficiency.

## Results

### Design and Functional Characterization of FL711

To generate an mRNA capable of producing a pharmacologically feasible TLR5 agonist, we designed FL711, an mRNA containing a mammalian codon-optimized open reading frame encoding a polypeptide with predicted TLR5-agonistic activity. This polypeptide is a derivative of bacterial flagellin extensively modified to (i) ensure functionality when translated in mammalian cells, (ii) enable extracellular secretion for extrinsic immunostimulation, and (iii) achieve substantial deimmunization to avoid neutralization by pre-existing antibodies. These goals were accomplished by mutating two glycosylation sites known to interfere with TLR5 agonist activity in mammalian cells (*27*), adding an N-terminal signal peptide, and removing – through multiple deletions and point mutations – major human T- and B-cell epitopes while preserving the core TLR5-binding D1 domain, as previously described (*26*) (**Fig. 1A**).

**Figure 1.**
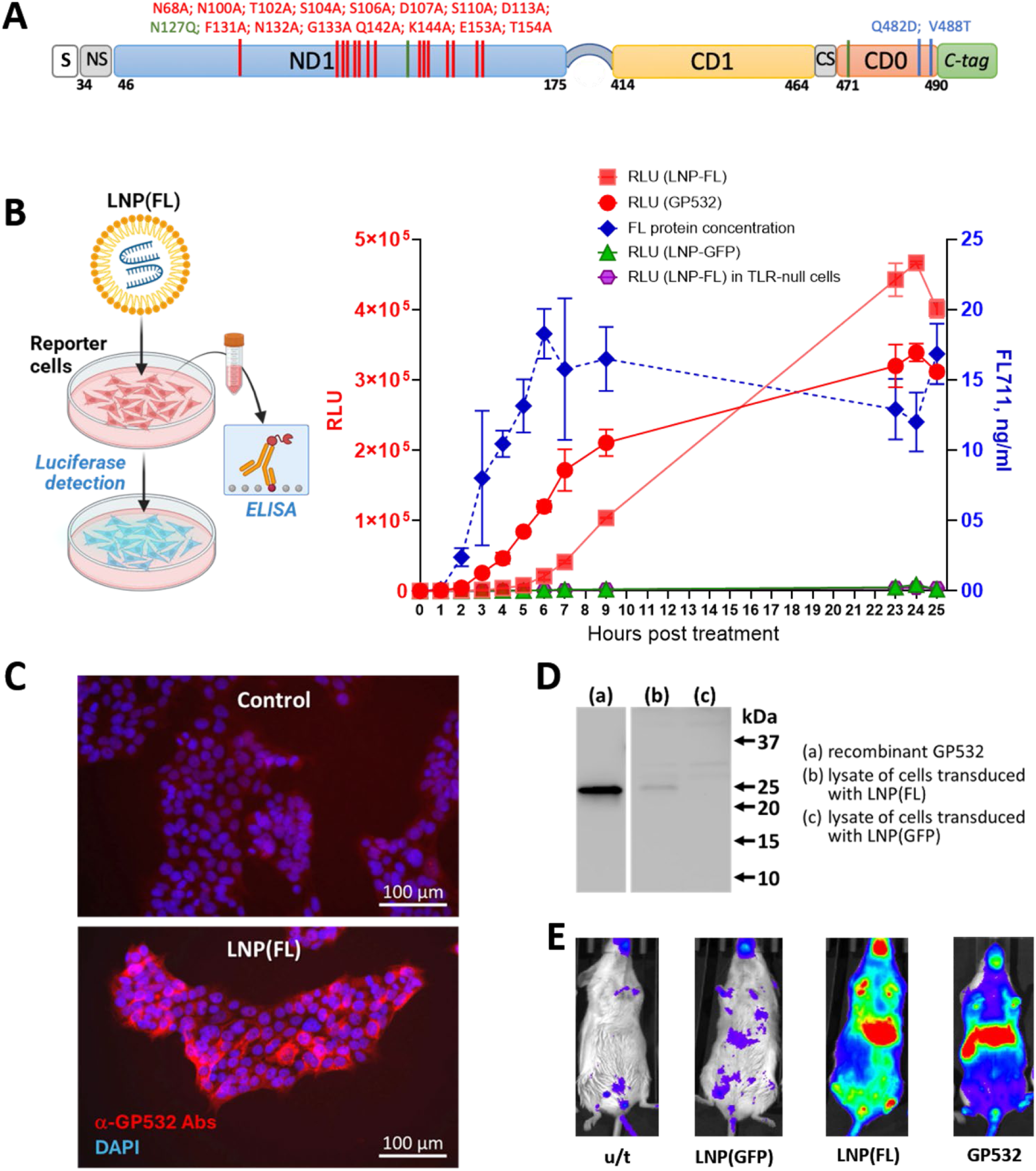
LNP-delivered FL711 mRNA drives TLR5-agonistic protein translation. **(A)** Schematic structure of FL711. Diagram illustrating the design of FL711 protein encoded by LNP-delivered FL711 mRNA encoding a truncated TLR5 agonist optimized for secretion and TLR5 activation. The construct includes an N-terminal secretion signal, modified flagellin-derived domains, and a C-terminal tag. **(B)** NF-κB activation and protein secretion kinetics. HEK293-TLR5-Luc reporter cells were treated with GP532 (50 ng/ml) or transfected with LNP-encapsulated mRNAs encoding FL711 or encoding green fluorescent protein (GFP). NF-κB–dependent luciferase activity (RLU, left Y-axis) and FL711 protein concentration in culture supernatant (ng/ml, right Y-axis) were measured up to 25 h post-treatment. HEK293-TLR-null reporter cells treated with LNP(FL711) served as a negative control for TLR5-independent NF-κB activation. Data represent mean ± SD of triplicates. **(C)** Immunofluorescence detection of FL711 protein. Cells were fixed at 24 h post-transfection and stained with anti-GP532 antibodies (red) and DAPI (blue). Strong cytoplasmic staining indicates cellular expression of FL711 protein after LNP(FL711) delivery. **(D)** Western blot confirmation of FL711 expression. Cell lysates collected 24 h after transduction with LNP(FL711) were analyzed by Western blot using anti-GP532 antibodies. A band corresponding to FL711-encoded protein (∼25 kDa) was detected, while no signal was observed in negative control (lysate of cells transduced with LNP(mRNA-GFP)). Recombinant GP532 protein was used as a positive control. (a) recombinant GP532; (b) lysate of cells transduced with LNP(FL711); (c) lysate of cells transduced with LNP(mRNA-GFP). **(E)** In vivo activation of the NF-κB signaling pathway in BALB/c-Tg(IκBα-luc)Xen mice following subcutaneous administration of LNP(FL711), LNP(mRNA-GFP), recombinant GP532. Representative in vivo bioluminescence images acquired 5 hours after intramuscular injection of 1 µg LNP(FL711), or 1 µg of non-coding LNP(mRNA-GFP) as a non-TLR5 agonistic RNA control, or recombinant protein GP532 (2 µg) as a positive control.

Synthetic FL711 mRNA (containing 100% N1-methylpseudouridine) was encapsulated in lipid nanoparticles (LNPs) formulated using a Moderna-like composition (see *Materials and Methods*) and subjected to functional testing (**Fig. 1B**). HEK293-hTLR5-Luc cells expressing TLR5, HEK293-hTLR4-Luc cells expressing TLR4, and HEK293-null-Luc cells lacking TLRs – all carrying an NF-κB–responsive luciferase reporter – were treated with LNP(FL711), and luciferase activity was measured at multiple time points following treatment . Recombinant TLR5 agonist GP532 – with structure closely resembling FL711-encodced polypeptide (*26*) – served as a positive control. A strong, time-dependent induction of luciferase, approaching the magnitude of activation produced by recombinant protein but with delayed onset, was observed in HEK293-hTLR5-Luc cells, but not in TLR5-negative reporter lines, confirming TLR5-specific NF-κB activation by the FL711-encoded product.

ELISA of supernatants from HEK293-null-Luc cultures showed gradual accumulation of secreted FL711 protein, mirroring the NF-κB activation profile (**Fig. 1B**, dashed blue line). Immunofluorescent staining with antibodies against TLR5 agonist GP532 showed intracellular signal in LNP(FL711)-transfected HEK293 cells, consistent with transient localization of flagellin derivative within the secretory pathway; control cells lacked specific staining (**Fig. 1C**). Western blot analysis confirmed expression of a full-length FL711 derivative of the expected size (∼25 kDa), matching the electrophoretic mobility of purified recombinant GP532 (**Fig. 1D**).

To assess LNP(FL711) functionality in vivo, we used BALB/c-Tg(IκBα-luc)Xen mice (*2*) expressing an NF-κB–responsive luciferase reporter, enabling real-time monitoring of NF-κB activation. Mice received intramuscular injections of LNP formulations of FL711 or mRNA encoding green fluorescent protein (mRNA-GFP, a non-TLR5-agonistic mRNA control) – both at dose of 1 μg of RNA. Recombinant TLR5 agonist GP532 (2 μg) served as a positive control; untreated animals served as baseline controls. As shown in **Fig. 1E**, LNP(FL711) induced luciferase expression, indicative of NF-κB activation, comparable in magnitude and anatomical distribution to recombinant TLR5 agonist, whereas LNP(mRNA-GFP) did not produce a detectable signal.

Together, these findings demonstrate that LNP(FL711) functions as an efficient producer of a secreted, biologically active TLR5 agonist capable of eliciting robust TLR5-dependent NF-κB activation *in vitro* and *in vivo*.

### Adjuvant activity of FL711 in the context of an anti-SARS-CoV-2 mRNA vaccine

To validate FL711 as an adjuvant for LNP(mRNA) vaccination, we used an experimental anti–SARS-CoV-2 mRNA vaccine consisting of LNP-formulated mRNA encoding the SARS-CoV-2 Spike protein (WA1, mammalian codon optimized, and containing HexaPro stabilizing mutations) (*28*). K18-hACE2 mice were vaccinated in a prime–boost regimen with a 21-day interval using LNPs containing 0.3 μg of Spike mRNA admixed with LNPs containing 0.01 μg of either mRNA-GFP or FL711 (**Fig. 2A**). The FL711 dose was selected as the lowest among the tested doses that demonstrated reliable immunization-enhancing activity in preliminary studies. (**Suppl. Fig. S1**).

**Figure 2.**
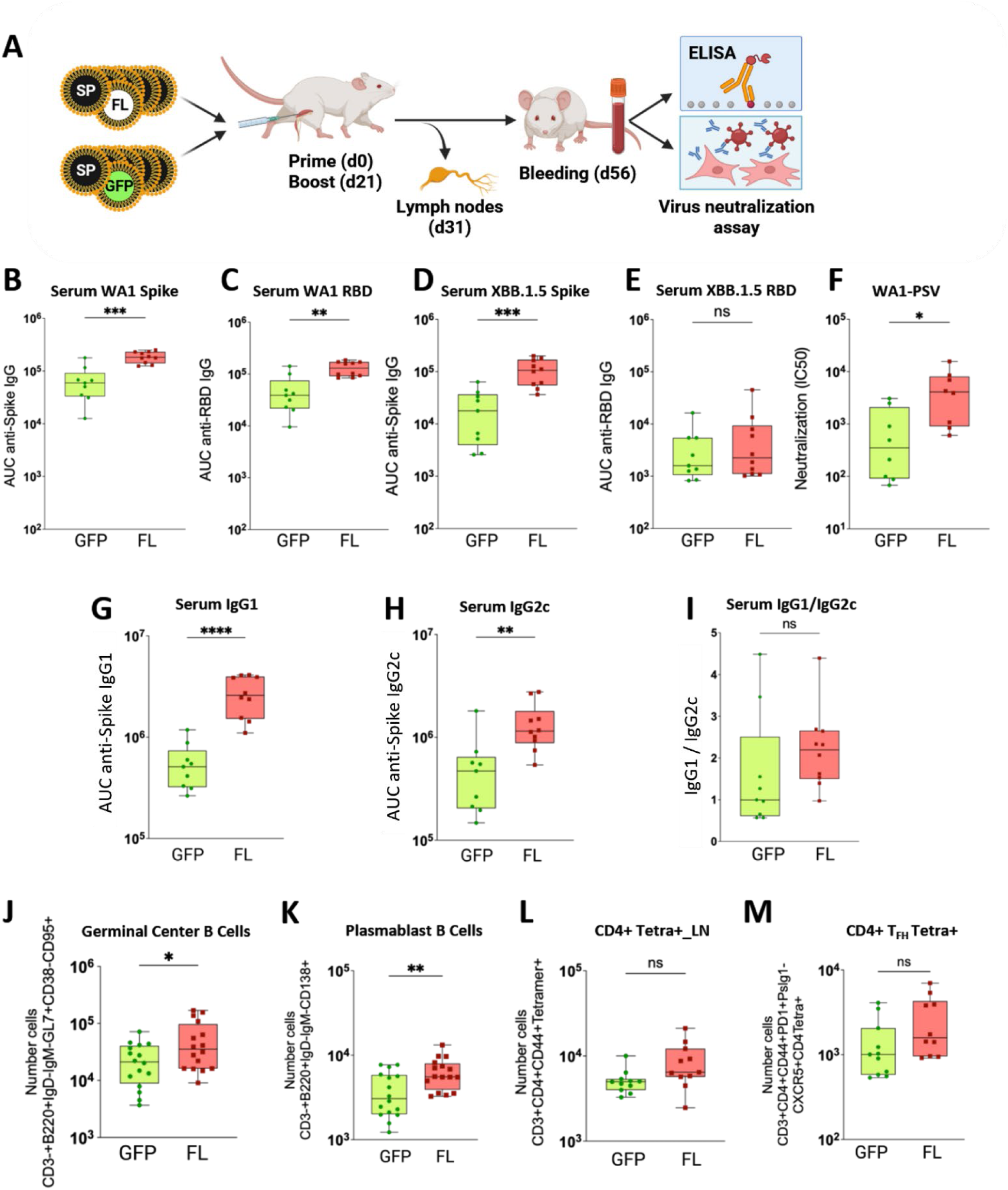
Co-delivery of FL711 mRNA improves humoral responses to SARS-CoV-2 Spike mRNA vaccination. **(A)** Scheme of the experiments. K18-hACE2 mice were intramuscularly (IM) immunized (primed) with either 0.3 μg of mRNA-LNPs encoding SARS-CoV-2 Spike protein combined with 0.01 μg of mRNA-LNPs encoding GFP or 0.3 μg of mRNA-LNPs encoding SARS-CoV-2 Spike protein combined with 0.01 μg of mRNA-LNPs encoding FL711 (FL). Mice received a booster immunization 21 days later with the same respective formulations. Serum was collected on day 56 post-prime (35 post-boost) to assess binding antibody responses against (**B)** SARS-CoV-2 WA1 full Spike IgG, **(C)** SARS-CoV-2 WA1 RBD–specific IgG, **(D)** SARS-CoV-2 XBB.1.5 Full Spike IgG, and **(E)** SARS-CoV-2 XBB.1.5 RBD–specific IgG. (**F**) Neutralizing activity was measured as serum IC50 against SCV2 WA1 Spike-pseudotyped vesicular stomatitis virus (VSV) in serum. IgG subclasses were also quantified: **(G)** IgG1, **(H)** IgG2c, and **(I)** the IgG1/IgG2c ratio. Boxes indicate median and interquartile range; whiskers indicate min–max; each point represents one mouse, and results are pooled from two independent experiments. Statistical significance was determined using Mann-Whitney U test; *P ≤ 0.05, **P ≤ 0.01, ***P ≤ 0.001, ****P ≤ 0.0001. **(J)** On day 31 post-prime (10 post boost), mice were euthanized, and inguinal lymph nodes (iLNs) were collected. Single-cell suspensions were prepared, stained, and analyzed to determine the absolute numbers of germinal center B cells, **(K)** plasmablast B cells, **(L)** CD4^+^ tetramer–positive T cells, and **(M)** CD4^+^ tetramer–positive follicular helper T cells. Boxes indicate median and interquartile range; whiskers show the full range; each point represents an individual mouse. Statistical significance was assessed using unpaired two-tailed t-test or Welch’s t-test, as appropriate. ns, not significant; *P ≤ 0.05; **P ≤ 0.01. Each data point represents an individual mouse, and results are pooled from two to three independent experiments.

ELISA and neutralization assays performed on serum collected at day 56 (35 days post-boost) showed a significant increase in binding IgG against homologous SARS-CoV-2 WA1 antigens (full Spike and RBD) (**Fig. 2 B,C**). These results confirm the adjuvant activity of FL711 for mRNA vaccines and that it is independent of additional LNP dose. Furthermore, ELISA performed using divergent XBB.1.5 Spike and RBD antigens revealed that FL711 enhanced the development of XBB.1.5 Spike-binding antibodies, but not XBB.1.5 RBD-binding antibodies, suggesting that this enhanced breadth of antibody reactivity targets conserved rather than variant-specific Spike epitopes (**Fig. 2D,E**). Importantly, consistent with increases in binding antibody titers, neutralizing antibody titers significantly increased by more than 10-fold (**Fig. 2F**). Corresponding to increased overall anti-SARS-CoV-2 Spike antibody levels, both IgG1 and IgG2c were increased in FL711 relative to the control. Additionally, there was a non-significant increase in the IgG1/IgG2 ratio, suggestive of a potential balancing of the Th1 and Th2 responses or skewing toward Th2 from the Th1-dominated mRNA-LNP response (**Fig. 2 G-I**).

Secondary lymphoid organs, such as lymph nodes and spleen, are essential for generating potent and durable antibody responses. To identify mechanisms by which FL711 enhances mRNA-LNP vaccine responses, we vaccinated mice as above and performed flow cytometry on draining lymph nodes 10 days post-boost (**Supplementary Fig. S2**). We observed a significant increase in plasmablast B cells – the precursors of long-lived plasma cells – and an increase in germinal center B cells (**Fig. 2J,K**). We also observed non-significant increases in antigen-specific total CD4⁺ T cells (MHC-II tetramer S62–76 (VTWFHAIHVSGTNGT)+) and antigen-specific CD4⁺ T follicular helper (T_FH_) cells (**Fig. 2L,M**). Together, these data suggest that enhanced activation of draining lymph nodes in the presence of FL711 contributes to improved antibody responses.

To assess whether FL711 enhances protective efficacy, vaccinated K18-hACE2 mice were challenged with 5 × 10^4^ PFU homologous (WA1) or divergent (XBB.1.5) SARS-CoV-2 at 41 days post-boost. Lung viral titers measured at 3 days post-infection showed significantly reduced viral loads in animals receiving FL711 adjuvant (3.6-fold median reduction for WA1 and 10-fold for XBB.1.5) relative to mRNA-GFP controls (**Fig. 3 B,C**).

**Figure 3.**
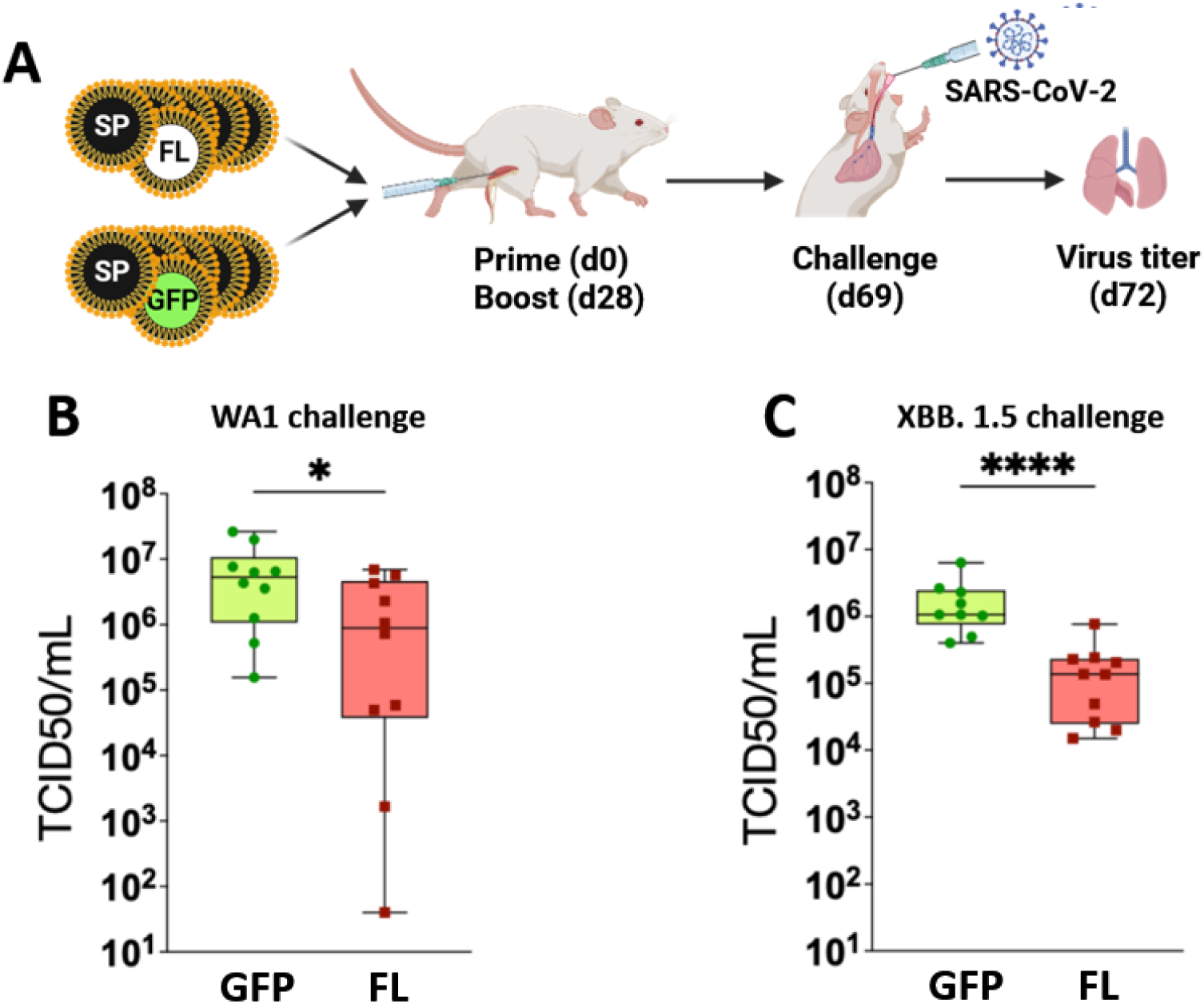
FL711 mRNA supplementation reduces lung viral titers after SARS-CoV-2 challenge. **(A)** Scheme of the experiments. K18-hACE2 mice were primed IM with either 0.3 μg of mRNA-LNPs encoding full-length SARS-CoV-2 Spike protein combined with 0.01 μg of mRNA-LNPs encoding GFP or 0.3 μg of mRNA-LNPs encoding full-length SARS-CoV-2 Spike protein combined with 0.01 μg of mRNA-LNPs encoding the FL711. Mice received an intramuscular boost with the same doses and formulations on day 21 post-prime. On day 62 post-prime (41 post-boost), mice were infected intranasally (IN) with 5 × 10⁴ plaque-forming units (PFU) of either SARS-CoV-2 **(B)** WA1 or **(C)** XBB.1.5 virus strains. On day 65 post-prime (3 post-infection), lungs were collected and TCID₅₀ assays were performed to determine viral titers. Boxes indicate median and interquartile range; whiskers show the full range; each point represents an individual mouse, and results are pooled from two independent experiments. Statistical significance was determined using the Mann-Whitney U test; *P ≤ 0.05, ****P ≤ 0.0001.

To evaluate whether FL711 enhances durability of immunity, we vaccinated mice as above and assessed antibody and neutralization titers at 103 days post-boost. We observed significantly improved durability of antibody responses, reflected by increased WA1 pseudovirus neutralization titers and increased serum IgG binding to XBB.1.5 Spike (**Fig. 4B,C**). Spike-binding antibody titers, while WA1 Spike-binding titers remained similar between control (GFP) and FL711 (**Fig. 4A**). At 117 days post-boost, mice were infected with XBB.1.5 SARS-CoV-2, and lung viral titers were measured at 3 days post-infection; weight loss and survival were monitored for 14 days (**Fig. 4D**). Mice receiving FL711–adjuvanted LNP (mRNA-Spike) exhibited improved protection from morbidity (**Fig. 4F**) and mortality (**Fig. 4E**) from heterologous challenge and significantly reduced lung viral loads at 3 days post infection compared to control animals that received the dose-matched LNP(mRNA-GFP).

**Figure 4.**
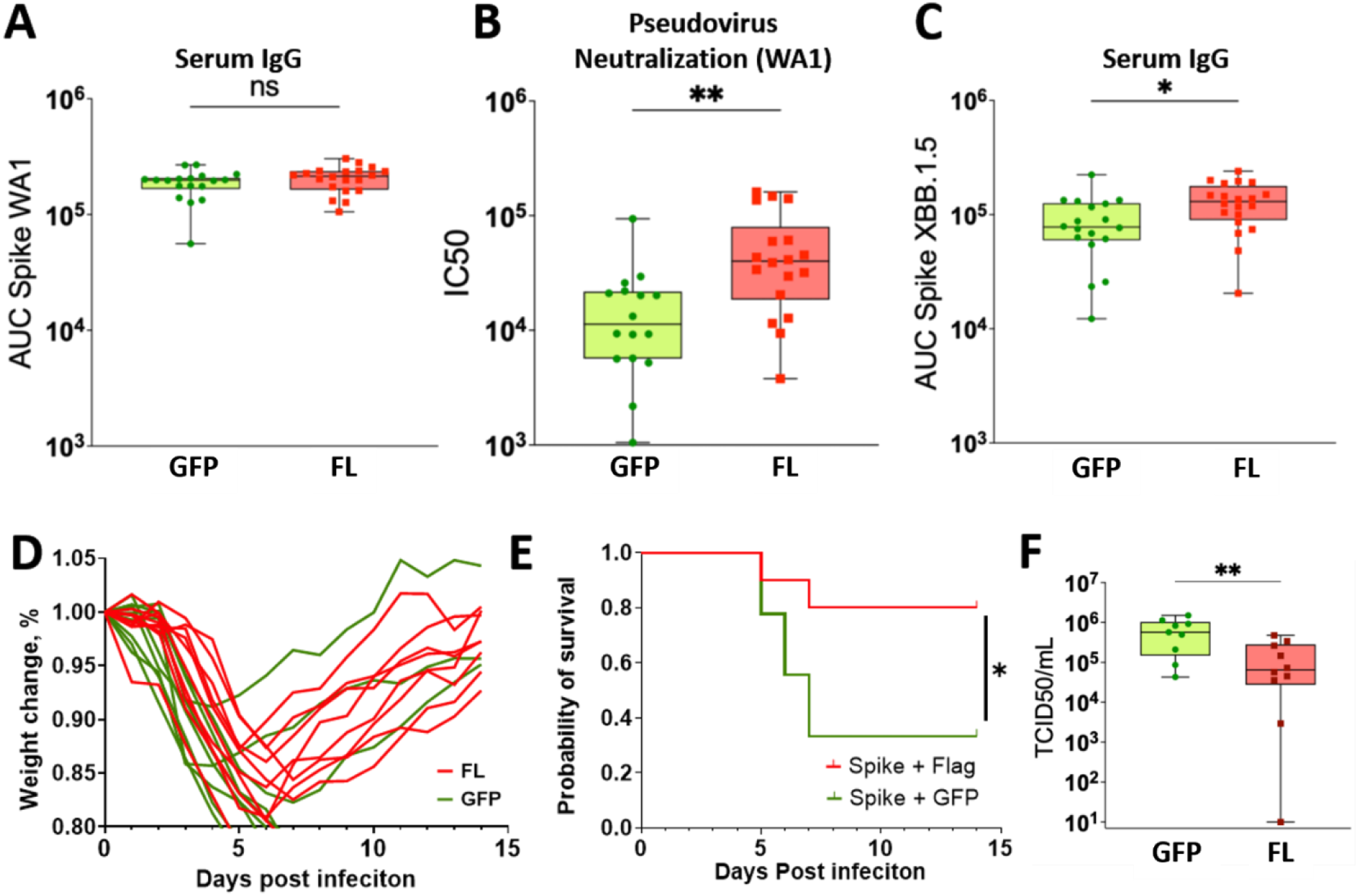
LNP(FL711) supplementation of anti-SARS-CoV-2 LNP(mRNA-Spike) vaccine promotes long-lasting humoral responses and durable protection against SARS-CoV-2. K18-hACE2 mice were primed intramuscularly with either 0.3 μg of LNP-encapsulated mRNAs encoding full-length SARS-CoV-2 Spike protein combined with 0.01 μg of mRNA encoding GFP or 0.3 μg of mRNA encoding full-length SARS-CoV-2 Spike protein combined with 0.01 μg of FL711. Mice were boosted intramuscularly with the same doses and formulations on day 21 post-prime. On day 125 post-prime (104 post-boost), sera were collected to assess binding antibody responses against **(A)** IgG SARS-CoV-2 WA1 Spike and **(B)** neutralization of SARS-CoV-2 WA1 Spike–pseudotyped vesicular stomatitis virus (VSV), as well as **(C)** IgG SARS-CoV-2 XBB.1.5 Spike. On day 138 post-prime (day 117 post-boost), mice were challenged intranasally with 5 × 10⁴ PFU of SARS-CoV-2 XBB.1.5. Mice were divided into two cohorts: one monitored for **(D)** weight loss over 14 days and **(E)** survival probability determined, and **(F)** a second cohort euthanized 3 days post-infection for viral load assessment by TCID₅₀. Each data point represents an individual mouse, and results are pooled from two independent experiments. Boxes indicate median and interquartile range; whiskers show the full range; each point represents an individual mouse, and results are pooled from two independent experiments. Statistical significance was determined using the Mann-Whitney U test, except for **(E)** where statistical significance was calculated by means of log-rank Mantel-Cox test; ns, not significant; *P ≤ 0.05; **P ≤ 0.01.

Taken together, these data provide orthogonal evidence supporting the use of FL711 as an effective mRNA-LNP vaccine adjuvant across multiple viral strains, enhancing both the breadth and durability of antibody responses and improving protection against viral infection.

### Adjuvant activity of FL711 in the context of an anti-influenza mRNA vaccine

We next evaluated the adjuvant activity of FL711 *in vivo* using an influenza mRNA vaccine consisting of LNP-formulated, synthetic pseudouridine-modified mRNA encoding the full-length hemagglutinin (HA) of influenza A/California/07/2009 (H1N1). BALB/c and C57BL/6 mice were tested using a standard prime-boost protocol. In all experiments, animals received intramuscular injections on days 0 and 14 of 0.5 μg of LNP(mRNA-HA) with or without co-administration of LNP(FL711), and antibody responses were assessed on day 51.

Functional neutralization of influenza virus by vaccine-induced antibodies was quantified using a microneutralization assay. The endpoint titer was defined as the reciprocal of the highest serum dilution achieving 100% neutralization (zero NP signal in the NP sandwich immunoassay). A benchmark of 1:40 was used as the putative protective threshold. As shown in **Fig. 5A,B**, the addition of FL711 increased vaccine efficacy in both mouse strains, as reflected by higher virus-neutralizing antibody titers. While the entire dose range of FL711 tested (0.03–0.3 µg) enhanced HA-binding IgG titers (**Suppl. Fig. S3**), the 0.3 µg dose was selected for subsequent experiments based on its superior improvement of virus-neutralizing antibody capacity.

**Figure 5.**
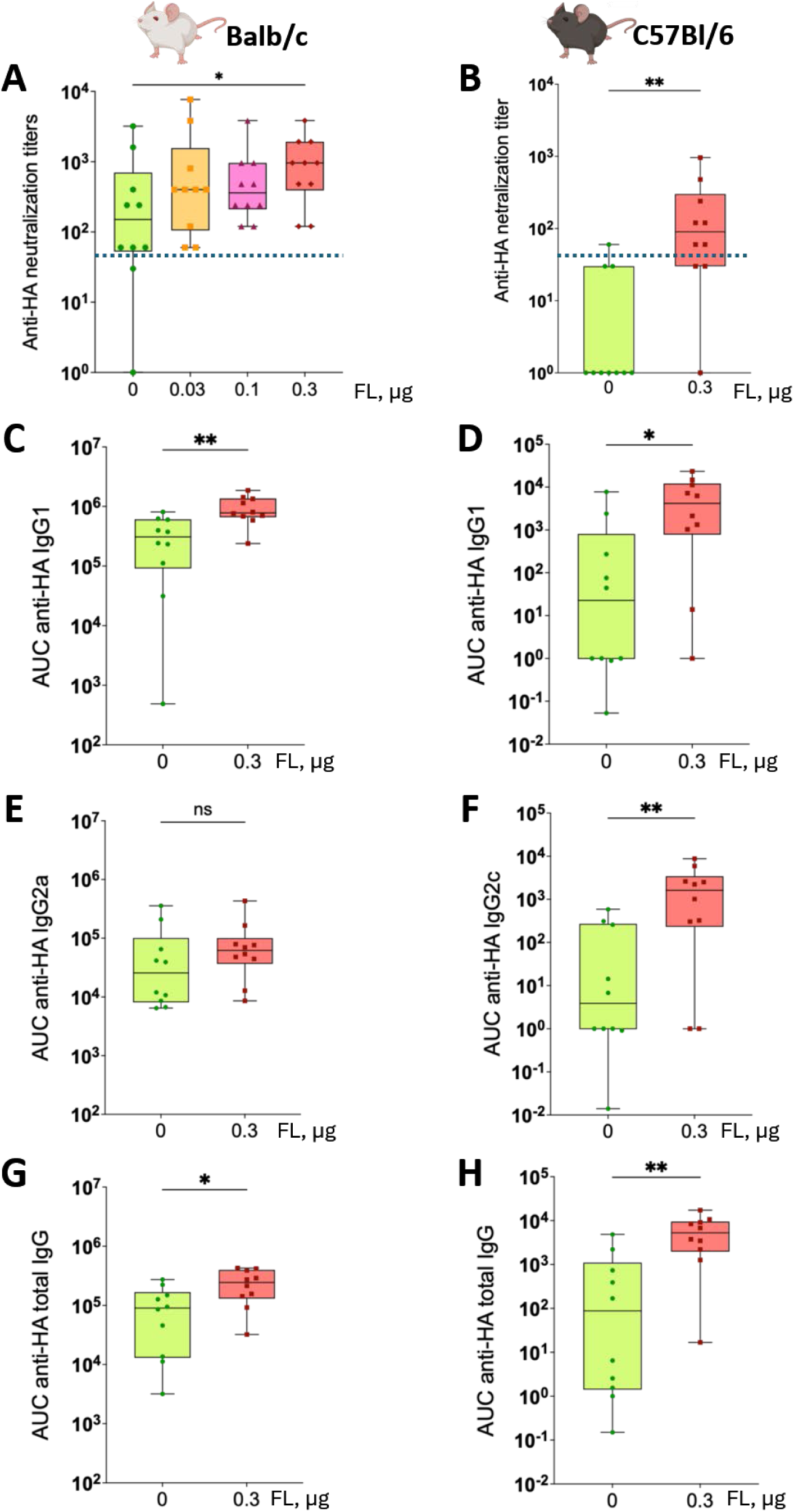
FL711 enhances neutralizing and HA-specific IgG subclass responses to influenza A mRNA vaccination in C57BL/6j and BALB/c mice. C57BL/6j and BALB/c mice were immunized intramuscularly on days 0 and 14 with 0.5 µg of LNP(HA), administered with or without LNP(FL711). Serum was collected on day 51 post-prime. **(A)** Anti-HA neutralizing antibody titers in BALB/c mice vaccinated with increasing doses of LNP(FL711): 0, 0.03, 0.1, or 0.3 µg. **(B)** Anti-HA neutralizing antibody titers in C57BL/6j mice vaccinated with LNP(mRNA-HA) alone or with 0.3 µg LNP(FL711). The dashed blue line denotes the protective threshold (titer = 40). **(C)** HA-specific IgG subclass responses in BALB/c mice, shown as AUC values for IgG1, IgG2a, and total IgG. **(D)** HA-specific IgG subclass responses in C57BL/6j mice, shown as AUC values for IgG1, IgG2c, and total IgG. Boxes indicate median and interquartile range; whiskers show the full range; each point represents an individual mouse. Statistical significance was determined using the Mann–Whitney test. ns, not significant; *P < 0.05; **P < 0.01.

Overall antibody responses in BALB/c mice were higher than those in C57BL/6 mice across all treatment groups, consistent with known strain-dependent immunological backgrounds: BALB/c mice are generally more prone to Th2-type humoral responses, whereas C57BL/6 mice mount more Th1-biased, cell-mediated immunity (*29*). The adjuvant effect of FL711 was evident in both strains; however, it was more pronounced in C57BL/6 mice, which are typically weaker responders. In C57BL/6 mice, addition of FL711 converted a largely subeffective dose of mRNA-HA into an effective one, with a greater fraction of animals surpassing the protective threshold (**Fig. 5B**). In BALB/c mice, the same vaccine dose was already substantially effective, leaving less room for improvement (**Fig. 5A**). The fact that enhancement was most pronounced at the lower, subeffective antigen dose demonstrates that FL711 can improve immunogenicity under antigen-sparing conditions, enabling reduction of antigen-encoding mRNA per vaccine dose without loss of efficacy.

IgG2a and IgG2c represent functionally analogous Th1-associated subclasses in mice but differ by strain genetics: BALB/c mice express IgG2a, whereas C57BL/6 mice lack IgG2a and instead express IgG2c (*30*). Accordingly, subclass-specific assays used IgG2a detection for BALB/c samples and IgG2c detection for C57BL/6 samples. Consistent with the results obtained with anti-SARS-CoV-2 vaccine (**Fig. 2)**, LNP(FL711) enhanced both Th2-associated IgG1 response and the strain-specific Th1-associated IgG subclasses (**Fig. 5C–F**), indicating that TLR5-agonistic mRNA amplifies the anti-HA antibody response without selectively skewing the subclass distribution toward either a Th1- or Th2-associated pattern.

These results show that the TLR5-agonistic LNP(FL711) acts as an effective adjuvant also for anti-influenza mRNA vaccine, enhancing functional antibody responses in both BALB/c and C57BL/6 mice and supporting a balanced Th1/Th2 response while enabling robust immunization even at sub-effective antigen-encoding mRNA doses.

### Biological effects of FL711 assessed by local and distal transcriptional responses

Effective vaccine adjuvants require the induction of an inflammatory response at the site of antigenic signal delivery. For mRNA vaccines, this signal arises from both components – synthetic mRNA and the LNP carrier – each capable of triggering innate immune receptors and thereby contributing to adjuvant activity. Response to synthetic RNA is mediated by several innate immune receptors, including TLR3, TLR7/8, PKR, RIG-I, and MDA5, which predominantly induce type I interferon (IFN-I) responses. Although activation of IFN-I response has adjuvant properties, it can also inhibit translation of the vaccine-encoded antigen from mRNA-based vaccines, thereby limiting immunogenicity (*31*). We hypothesized that enhancing NF-κB activation – the principal pathway triggered by TLR5 – while minimally activating IFN-I would improve and balance the overall immune response to mRNA-LNP vaccination. Additionally, TLR5 has not been implicated in the adjuvant response mediated by commercial LNP formulations, suggesting that, by activating TLR5, we may achieve a synergistic effect with the existing adjuvant properties of the mRNA and the LNP. As shown in previous sections, this strategy was effective. Here, we sought to define the specific immunological programs contributed by TLR5-agonistic mRNA in the context of LNP(mRNA) vaccination. The experimental design is shown in **Fig. 6A**.

**Figure 6.**
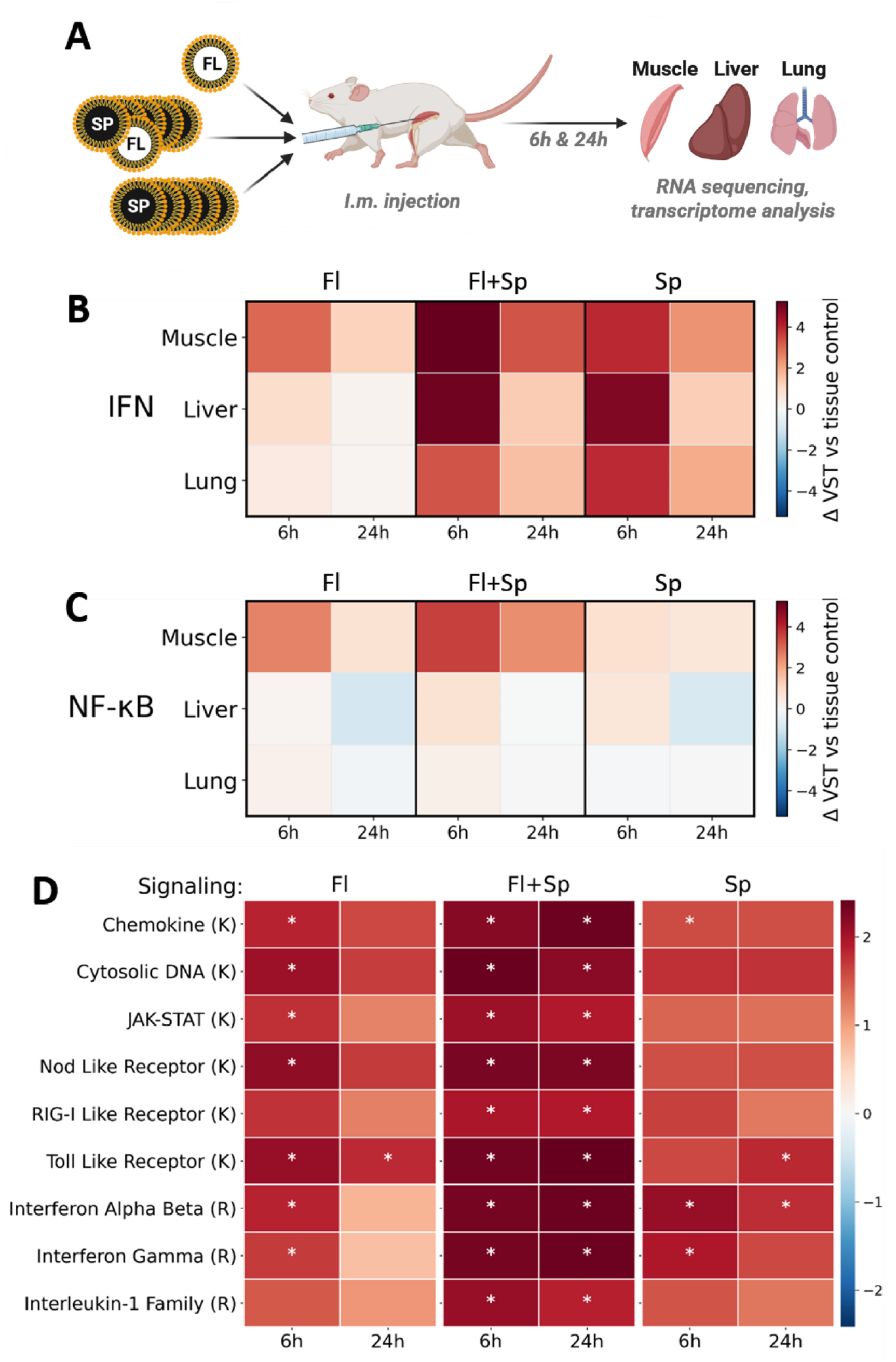
The effects of FL711 supplement to anti-SARS-CoV-2 mRNA vaccine on the local and systemic transcriptional response. **(A)** Experimental design. C57BL/6 female mice (9 weeks-old, n = 3 per group) received IM injections of LNP(mRNA) formulations containing 0.1 μg of FL711 mRNA (Fl), 0.5 μg of Spike mRNA (Sp), or both FL711 (0.1 μg) and Spike (0.5 μg) mRNAs (Fl+Sp). Injected muscle, liver, and lungs were collected at 6 h and 24 h post injection. RNA was isolated, sequences and transcriptomes were analyzed comparing them to those from the organs of age- and sex-matched untreated control mice. **(B)** IFN-I-associated transcriptional response across tissues and timepoints. Heatmap values show mean ΔVST expression relative to tissue-matched untreated controls for predefined IFN-I-responsive genes. **(C)** NF-κB-associated transcriptional response across tissues and timepoints. Heatmap values show mean ΔVST expression relative to tissue-matched untreated controls for predefined NF-κB-responsive genes. **(D)** Enrichment analysis of inflammatory and innate-immune signaling pathways in muscle. Preranked GSEA was performed using tissue- and time-matched descriptive log2FC-ranked gene lists. Color indicates fgsea normalized enrichment score (NES); asterisks indicate FDR-adjusted fgsea pathway-enrichment p-values. K and R denote KEGG and Reactome pathways, respectively.

C57BL/6 mice were injected intramuscularly with LNP(mRNA) formulations containing FL711 mRNA (0.1 μg per injection), mRNA encoding SARS-CoV-2 Spike protein (0.5 μg per injection), or their combination. At 6 and 24 hours post-injection, mice (n=3 per group) were sacrificed, and RNA was isolated from injected muscle as well as from liver and lung. Pooled RNA samples were sequenced and RNA-seq profiles were compared with tissue-matched untreated controls to define treatment-associated transcriptional shifts. Liver and lung were selected as distal organs known to respond to TLR5 agonists (*32*), (*26*), enabling assessment of local versus systemic effects of the vaccine with or without TLR5-agonistic mRNA. Results are shown in **Fig. 6B–D**.

As expected, all vaccine compositions induced a strong local and weaker distal IFN-I response, with magnitude roughly proportional to the mRNA dose (**Fig. 6B**). Addition of FL711 prolonged the IFN-I response, as indicated by a less pronounced decline in IFN-responsive transcripts at 24 hours compared with preparations containing each mRNA alone. Distal responses in lung and liver were minimal following LNP(FL711) alone, indicating that this dose (0.1 μg) did not generate circulating IFN-α/β at levels sufficient to elicit systemic transcriptional effects. Expression patterns of individual IFN-responsive genes are shown in **Supplementary Fig. S3A.**

Gene-level expression profiles and pathway summaries highlighted NF-κB-associated inflammatory programs as prominent transcriptional changes linked to FL711 supplementation (**Fig. 6C**), with individual NF-κB-responsive transcripts shown in **Supplementary Fig. S4B**. NF-κB activation was substantially weaker in distal organs (with the exception of several transcripts such as *Il1b* and *Tlr2*). In contrast, NF-κB-mediated transcription in the injection site was stronger and more sustained when LNP(FL711) was combined with LNP(mRNA-Spike).

The enhancement of the local inflammatory response by FL711 is demonstrated by gene-set enrichment analysis (GSEA) of tissue- and time-matched expression contrasts using focused KEGG and Reactome pathway sets. Statistical assessment of pathway engagement using Fast Gene Set Enrichment Analysis (fgsea) normalized enrichment scores (NES), with FDR-adjusted enrichment p-values used to denote enriched pathways, showed a substantial increase in both the magnitude and duration of all major inflammatory signaling programs at the injection site (muscle) when LNP(mRNA-Spike) was supplemented with mRNA(FL711) (**Fig. 6D**). These findings provide a mechanistic explanation for the observed and previously described improvement in vaccine performance.

These observations align with the functional data demonstrating improved vaccine efficacy with FL711 supplementation. FL711 enhances immunogenicity by inducing a robust NF-κB response that complements the IFN-I signaling elicited by LNP(mRNA) and extends the duration of local inflammatory activation. The minimal systemic inflammatory signature produced by FL711 indicates that its adjuvant activity can be achieved without triggering systemic inflammation, supporting the favorable safety profile of this mRNA-encoded TLR5 agonist.

## Discussion

Antigen-encoding mRNA designs that leverage modified nucleosides, dsRNA structures, or optimized untranslated regions have been shown to tune innate immune sensing and enhance immunogenicity (*33*), (*34*). Compared with these approaches, mRNA-encoded protein adjuvants offer the unique advantage of spatially and temporally coordinated expression with the antigen, potentially improving efficacy while minimizing systemic exposure. While many current and previous studies have focused on incorporating host-derived factors such as cytokines or other signaling molecules, in this study, we demonstrate that supplementing LNP(mRNA) vaccines with a small proportion of FL711, a synthetic mRNA encoding a deimmunized secreted TLR5 agonist, substantially enhances immune responses across two distinct antiviral models – SARS-CoV-2 and influenza A.

FL711 should be viewed within the broader and rapidly evolving landscape of mRNA-encoded adjuvants. Several alternative mRNA-compatible adjuvant strategies have been explored, including co-delivery of mRNAs encoding cytokines such as IL-12 and GM-CSF (*18*), (*35*), (*36*), and mRNAs encoding costimulatory or innate-activating proteins such as CD40L, CD70, and constitutively active TLR4 in the TriMix platform (*37*). Notably, mRNA encoding constitutively active STING enhances antigen-specific T-cell immunity by promoting robust IFN-I signaling (*19*). In addition, noncoding immunostimulatory RNA adjuvants that activate TLR7/8 and RIG-I pathways, such as RNAdjuvant/CV8102, have also been developed (*38*). Continued comparative evaluation of these platforms will be important for defining optimal combinations and context-specific applications. TLR5 agonists occupy a distinct position among prospective mRNA-encoded adjuvants because of their unique combination of immunostimulatory and pharmacological properties, which together make them highly attractive candidates for vaccine enhancement (see *Introduction* for details). Interestingly, TLR5-mediated sensing of symbiotic microbial products has been identified as a key determinant of vaccine responses in humans and pre-clinical models (*39*),(*40*). The ability of TLR5 signaling to enhance vaccine response is clearly demonstrated in our experiments with FL711 mRNA, as discussed below.

A key finding of this work is that FL711 improves the qualitative features of the immune response. In both vaccine systems, co-delivery increased neutralizing antibody titers and expanded serologic breadth, including improved recognition of antigenically divergent SARS-CoV-2 variants. Mechanistically, TLR5-driven NF-κB activation complements the IFN-I and non-IFN-I innate signature induced by conventional LNP(mRNA) formulations, resulting in enhanced lymph node activation, increased GC-B cell, plasmablast, and antigen-specific CD4⁺ T cell responses, and a more balanced Th1/Th2 profile. This integrated innate signaling is consistent with prior work showing that combinatorial PRR activation can improve both humoral and cellular immunity (*41*), (*42*). The effects of FL711 were reproducible across two mouse strains and were achieved with minimal additional mRNA input, underscoring the potency of encoded TLR5 agonism as an adjuvant strategy (schematic interpretation of FL711 adjuvant activity is shown in **Fig. 7**).

**Figure 7.**
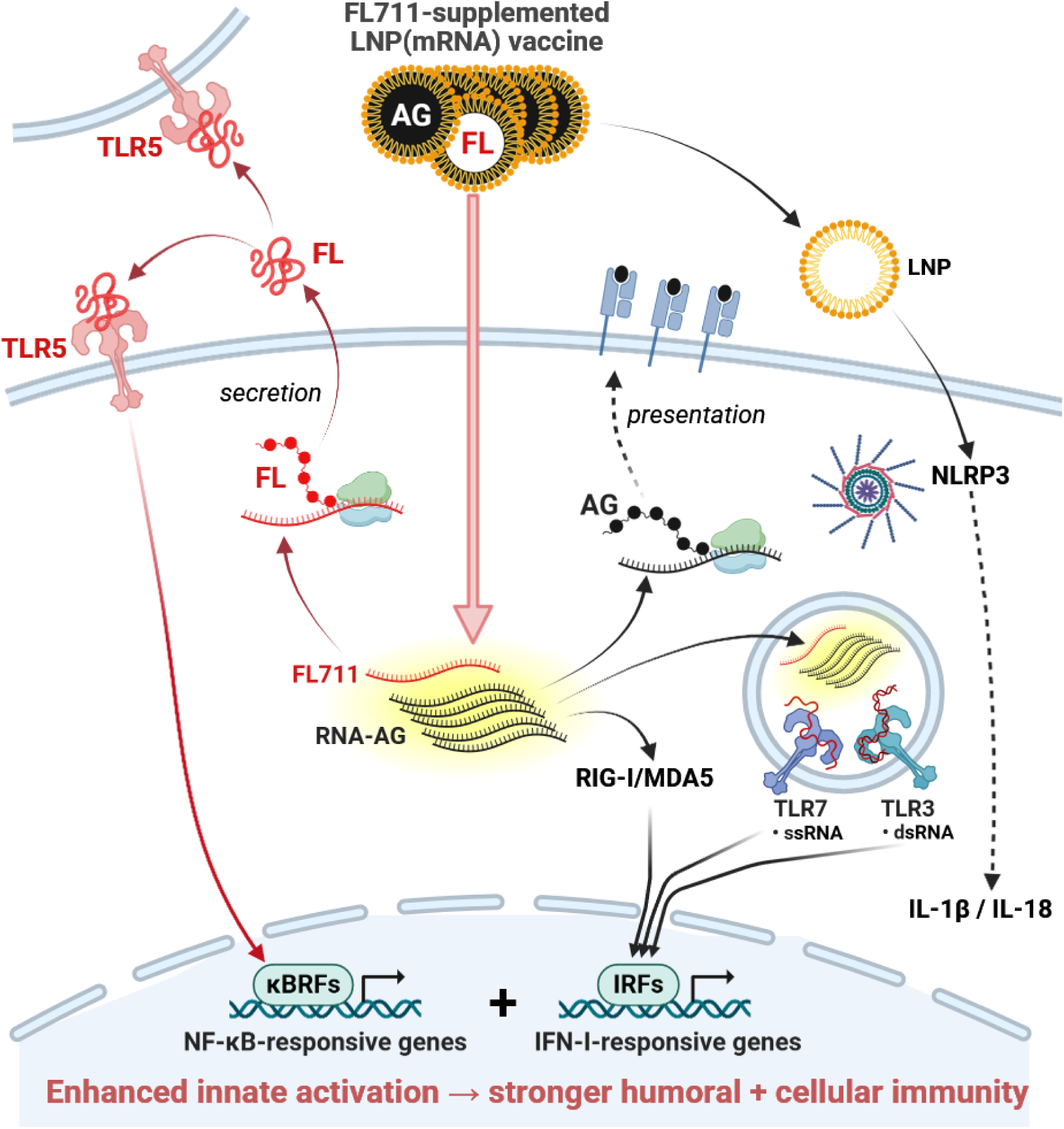
Proposed mechanism of the adjuvanting activity of FL711 in the LNP(mRNA) vaccine. The vaccine co-encapsulates antigen-encoding mRNA (AG) and FL711 (FL) within a single lipid nanoparticle (LNP). Upon cellular uptake, two parallel adjuvanting mechanisms are engaged. First, the LNP vehicle and its RNA cargo trigger innate immune sensing through complementary pathways: cytoplasmic RNA sensors (RIG-I/MDA5) and endosomal Toll-like receptors (TLR3, TLR7) activate interferon regulatory factors (IRFs), driving IFN-I–responsive gene expression; concurrently, the LNP itself activates the NLRP3 inflammasome, resulting in IL-1β and IL-18 secretion. Second, FL711 is translated into protein (FL), which activates TLR5 both in an autocrine and in a paracrine manner, driving NF-κB–responsive gene expression via κB-responsive transcription factors (κBRFs). In parallel, AG mRNA is translated and the resulting antigen is presented via surface MHC molecules to T cells. Together, TLR5-mediated NF-κB activation and LNP(mRNA)-intrinsic innate sensing converge to enhance both IFN-I and pro-inflammatory cytokine responses, resulting in stronger humoral and cellular immunity.

Beyond response quality, FL711 also extended the durability of immunity, with sustained neutralizing titers and improved protection upon delayed viral challenge. This likely reflects enhanced germinal center dynamics, supported by increased plasmablast formation and T follicular helper (Tfh) cell activity – both essential for long-lived antibody responses and affinity maturation (*43*). Importantly, transcriptional profiling indicates that while the doses of FL711 used here prolong local immune stimulation, they do not increase systemic inflammation, supporting a favorable prospective safety profile.

Another major observation is the ability of FL711 to enable antigen-sparing vaccination. In both SARS-CoV-2 and influenza models, the addition of FL711 converted suboptimal antigen doses into fully protective regimens. This property has important implications for manufacturing scalability and global vaccine deployment, particularly in pandemic settings where dose-sparing strategies can substantially expand supply (*44*).

A further advantage of FL711 lies in its modularity. As an mRNA-encoded adjuvant, it can be readily incorporated into existing LNP platforms without substantial changes to manufacturing workflows. This is particularly relevant for rapidly adaptable vaccine platforms, such as seasonal or pandemic influenza vaccines. Moreover, its ability to enhance T cell activation and support balanced helper responses favors applicability to cancer vaccines, where weak immunogenicity of tumor antigens remains a major limitation.

Importantly, TLR5 signaling appears relatively preserved with aging, in contrast to several innate immune pathways that decline with immunosenescence, and its stimulation has demonstrated geroprotective effects in mouse models (*45*). This feature, combined with prior clinical observations of tolerability and immunization-enhancing efficacy of TLR5 agonists in older animals and humans (*25*), suggests that FL711 may be particularly valuable for vaccines targeting elderly populations, a group with historically reduced responsiveness to vaccination; however, this will require further investigation.

The use of TLR5-agonistic mRNA enables harnessing an innate immune program that has been evolutionarily optimized to detect and counteract microbial invasion. This is reflected in the distinctive tissue distribution of TLR5, which is enriched at key anatomical “infection gates” and targets, including the liver, lung, and bladder (*32*), (*46*), (*47*). Consequently, this adjuvanting mechanism may be especially valuable in the context of mucosal vaccination, where epithelial TLR5 expression provides a unique opportunity for potent, localized immune activation (*32*), (*46*). This potential remains to be explored using alternative mRNA delivery platforms that target mucosal surfaces.

In summary, these results suggest that mRNA-encoded TLR5 agonists have the potential to reshape mRNA vaccine design across multiple applications: (i) next-generation anti-infective vaccines requiring enhanced breadth and durability; (ii) therapeutic cancer vaccines, where robust T cell priming is critical; and (iii) mucosal or intranasal vaccines. The preclinical efficacy signals and favorable safety profile observed here, together with prior clinical experience with recombinant TLR5 agonists, support the feasibility of early-phase clinical translation in the context of anti-infective or anti-cancer mRNA vaccines.

## Materials and Methods

### Study performance and experimental sites

The study was conducted across multiple institutional sites. In vitro reporter assays, influenza vaccination studies, in vivo NF-κB bioluminescence imaging, influenza serology, influenza microneutralization assays, and RNA-seq sample preparation were performed at Roswell Park Comprehensive Cancer Center. SARS-CoV-2 vaccination, challenge, serological, neutralization, viral-load, and flow-cytometry studies were performed at Yale University, as indicated below. All animal procedures were approved by the corresponding Institutional Animal Care and Use Committees (IACUC).

### mRNA synthesis and LNP formulation

Synthetic mRNAs encoding full-length influenza hemagglutinin from A/California/07/2009 (H1N1), SARS-CoV-2 Spike, GFP, or the engineered TLR5 agonist FL711 were generated by in vitro transcription by TriLink Biotechnologies (part of Maravai Life Sciences, San Diego, CA) and formulated into lipid nanoparticles by Particella Inc. The mRNAs contained a 5′ cap, 5′ and 3′ untranslated regions, an open reading frame encoding the indicated protein, and a poly(A) tail.

In vitro mRNA synthesis by TriLink uses T7 RNA polymerase involving designing a DNA template with a T7 promoter, which is transcribed into RNA. The DNA template can be amplified via PCR or synthesized directly. In an in vitro transcription reaction, the purified DNA template is mixed with T7 RNA polymerase, transcription buffer, and nucleotides (N1-methyl-pseudouridine substitution for uridine), and then incubated at 37°C to allow RNA synthesis.

LNPs were prepared using a Precision NanoAssemblr system (Precision NanoSystems, Vancouver, BC) with a previously described SM-102-based lipid composition consisting of SM-102, DSPC, cholesterol, and PEG2000-DMG at a molar ratio of 50:10:38.5:1.5 (*48*). The final formulation buffer contained 20 mM Tris, 70 mM NaCl, and 10% sucrose.

mRNA integrity was assessed by capillary electrophoresis using an Agilent Bioanalyzer. Particle size and polydispersity index were measured using a Malvern Zetasizer, and mRNA encapsulation efficiency was determined using the RiboGreen assay. LNP formulations were stored at −80°C, thawed on ice, and used immediately after thawing. Unless otherwise indicated, all LNP doses are reported as mRNA mass.

### Cell lines and culture conditions

HEK293-hTLR5-Luc, HEK293-hTLR4-Luc, and HEK293-null-Luc NF-κB luciferase reporter cell lines were used to assess receptor-specific NF-κB signaling in vitro. Parental HEK293-hTLR5, HEK293-hTLR4, and HEK293-null cells were obtained from InvivoGen. The corresponding luciferase reporter lines were generated by lentiviral transduction of parental cells with an NF-κB–responsive firefly luciferase reporter cassette. HEK293-null-Luc cells were derived from HEK293-null cells carrying a pUNO control plasmid and served as a receptor-negative NF-κB reporter control. HEK293-hTLR4-Luc cells were included as a TLR4-expressing reporter control to assess receptor specificity.

Vero E6 cells stably expressing human angiotensin-converting enzyme 2 (ACE2) and transmembrane protease serine 2 (TMPRSS2) were used for SARS-CoV-2 studies. These cells were obtained through BEI Resources, NIAID, NIH: Cercopithecus aethiops kidney epithelial cells expressing TMPRSS2 and human ACE2, NR-54970.

All cells were maintained at 37°C in a humidified incubator with 5% CO₂ and cultured in Dulbecco’s modified Eagle medium (DMEM; Thermo Fisher Scientific) supplemented with 10% fetal bovine serum (FBS; 10437-010, Gibco) and 1% penicillin–streptomycin (30-002-CV, Corning). Cell viability was routinely assessed using a NucleoCounter NC-200 (Chemometec).

### In vitro treatment of HEK293 reporter cells with LNP(FL711)

For time-course experiments, HEK293-hTLR5-Luc, HEK293-hTLR4-Luc, and HEK293-null-Luc cells were seeded in 96-well tissue culture plates (3610, Costar) at 2.5 × 10⁴ cells per well in 100 µL of complete DMEM and incubated overnight before treatment. Cells were then exposed for 1 hour at 37°C to lipid nanoparticle–encapsulated LNP(FL711) or control LNP(mRNA-GFP) at a final mRNA concentration of 50 ng/mL. Recombinant GP532 protein was included as a positive control at 50 ng/mL. After the 1-hour exposure, the treatment medium was removed and replaced with fresh complete DMEM. Cells were further incubated at 37°C and sampled at defined time points from 1 to 25 hours. Culture supernatants were collected to quantify secreted FL711 protein, and corresponding cell lysates were analyzed for NF-κB–dependent luciferase activity.

### NF-κB luciferase reporter assay

NF-κB–dependent luciferase activity was measured in HEK293 reporter cells following treatment with LNP(FL711), control LNP(mRNA-GFP), or recombinant GP532 protein (recombinant TLR5 agonist GP532 resulted from a prior published work of our team (*26*)). At the indicated time points, culture supernatants were collected for quantification of secreted FL711 protein, and cells were lysed for measurement of intracellular luciferase activity. To measure firefly luciferase activity 20 µL of lysis buffer containing 50 mM Tris, pH 7.8, 25 mM NaCl, 1 mM MgCl₂, 0.1 mM EDTA, 20 mM DTT, 0.25 mM ATP, 0.25 mM D-luciferin potassium salt, and 0.25% Triton X-100 was added per well and luminescence signal was measure after 1 min at room temperature using a SpectraMax M3 or Tecan Infinite M1000 plate reader, with an integration time of 1000 ms per well. Relative luminescence units were recorded. Data were analyzed using GraphPad Prism and are presented as RLU or normalized reporter activity, as indicated in the figure legends.

### FL711 protein ELISA

Secreted FL711 protein in culture supernatants was quantified using an in-house sandwich ELISA. High-binding 96-well plates (437111, Thermo Fisher Scientific) were coated overnight at 4°C with affinity-purified rabbit anti–GP532 polyclonal antibody (generated in-house) at 2 µg/mL in PBS (21-040-CV, Corning). Plates were washed with PBS containing 0.05% Tween 20 (P2287, Sigma-Aldrich) and blocked for 1 hour at room temperature with casein blocking buffer (37528 Thermo Fisher Scientific) containing 0.05% Tween 20. Culture supernatants and recombinant GP532 protein standards were added in triplicate and incubated for 1 hour at room temperature. Recombinant GP532 reference standard was serially diluted in complete DMEM to generate standard curves. After washing, plates were incubated for 1 hour with biotinylated goat anti–GP532 polyclonal antibody (generated in-house) diluted 1:2,000 in assay buffer, followed by streptavidin–HRP (DY998, R&D Systems) for 1 hour at room temperature. After washing, QuantaBlu fluorogenic peroxidase substrate (15169, Thermo Fisher Scientific) was added, and plates were incubated for 20 min at room temperature with gentle shaking. Fluorescence was measured at 320 nm excitation and 420 nm emission using a SpectraMax M3 or Tecan Infinite M1000 plate reader. FL711 protein concentrations in cell culture were interpolated from the standard curve. Samples above the upper limit of quantification were reanalyzed after dilution.

### Animals

BALB/c-Tg(IκBα-luc)Xen reporter mice, which carry a luciferase reporter gene under the control of the NF-κB–responsive IκBα promoter, were used for in vivo bioluminescence imaging of NF-κB activation. 8-12-week-old female C57BL/6 and BALB/c mice were used for influenza immunization studies. BALB/c-Tg(IκBα-luc)Xen mice were obtained from PerkinElmer (Waltham, MA, USA), and C57BL/6 and BALB/c mice were obtained from The Jackson Laboratory (Bar Harbor, ME, USA). Mice were maintained under specific pathogen–free conditions at the Roswell Park Comprehensive Cancer Center animal facility. All animal procedures were approved by the Roswell Park Comprehensive Cancer Center Institutional Animal Care and Use Committee.

Parental transgenic K18-hACE2 mice [B6.Cg-Tg(K18-ACE2)2Prlmn/J, stock no. 034860] were obtained from The Jackson Laboratory. Mice were mated, bred, and housed in the Yale University animal facilities. Male and female mice, 8–12 weeks old, were vaccinated and used for SARS-CoV-2 immunization and challenge studies. All animal procedures, including sex, cage, and age matching, complied with federal regulations and were approved by the Yale School of Medicine Institutional Animal Care and Use Committee. In accordance with institutional policy to minimize animal use, control cohorts were shared across experiments when appropriate, with such instances noted in the figure legends. Group sizes were determined empirically, guided by prior publications using comparable experimental models. No predetermined statistical methods were applied to establish sample size. Age- and sex-matched animals were randomly assigned to treatment groups.

### In vivo NF-κB bioluminescence imaging

In vivo NF-κB activation was assessed in BALB/c-Tg(IκBα-luc)Xen reporter mice by bioluminescence imaging. Mice were injected intramuscularly with 20 µL of the indicated formulations, and imaging was performed 3 hours after injection. For imaging, mice received 3 mg of D-luciferin potassium salt (Gold Biotechnology) by intraperitoneal injection. Five minutes after luciferin administration, animals were anesthetized with 2.5% isoflurane, and bioluminescence images were acquired using an IVIS Spectrum system (PerkinElmer) with a 10-s exposure time. Images were analyzed using Living Image software (Caliper Life Sciences). Regions of interest were drawn over the injection site, and bioluminescence intensity was quantified as total flux in photons per second.

### Influenza mRNA vaccination studies and serum collection

8-12-week-old female C57BL/6 and BALB/c mice were immunized intramuscularly in the quadriceps with LNP(mRNA) formulations on days 0 and 14 using 29½-gauge insulin syringes. All influenza antigen–containing groups received 0.5 µg of LNP(mRNA-HA) per mouse.

For C57BL/6 studies, mice received 0.5 µg of LNP(mRNA-HA) alone or in combination with 0.3 µg of LNP(FL711) per mouse. For BALB/c studies, mice received 0.5 µg of LNP(mRNA-HA) alone or in combination with 0.3, 0.1, or 0.03 µg of LNP(FL711) per mouse.

Blood samples were collected by submandibular bleeding on day 28 for BALB/c mice and by cardiac puncture on day 51 for C57BL/6 mice. Serum was separated by centrifugation at 2,000 × g for 10 min at 4°C and stored at −20°C until analysis.

### SARS-CoV-2 vaccination and challenge studies

K18-hACE2 mice were vaccinated intramuscularly in the left quadriceps with 0.3 µg of LNP(mRNA-Spike) encoding SARS-CoV-2 Spike, in combination with 0.01 µg of either LNP(FL711) or control LNP(mRNA-GFP), diluted in PBS to a final volume of 20 µL. Mice were boosted with the same doses 28 days after the initial vaccination. Prior to vaccination or boosting, mice were anesthetized according to the approved institutional protocol. The SARS-CoV-2 isolate hCoV-19/USA-WA1/2020 (NR-52281, BEI Resources, Lot: 70044427) and the SARS-CoV-2 isolate hCoV-19/USA/MD-HP40900/2022, lineage XBB.1.5; Omicron variant (NR-59104, BEI Resources, Lot: 70057837), were used for challenge studies. Virus stocks were propagated in Vero E6-ACE2/TMPRSS2 cells, and infectious titers were determined by plaque assay as previously described (*49*),(*50*). Vaccinated or unvaccinated mice were inoculated intranasally with 5 × 10⁴ plaque-forming units of the indicated SARS-CoV-2 strain on the indicated day post-vaccination. Animals were monitored daily for clinical signs of disease and weight loss. Lungs were collected at 3 days post-infection for challenge experiments, and mice were euthanized 14 days post-infection for weight-loss studies.

### Anti-HA antibody ELISA

Anti-HA antibody titers were measured in mouse sera by indirect ELISA. High-binding MaxiSorp (442404, Thermo Fisher) 96-well plates were coated overnight at 4°C with recombinant HA1 from influenza A/California/07/2009 (H1N1) (IT-003-SW12p, Immune Technology Corp.) at 0.6 µg/mL in PBS. Plates were washed with PBS containing 0.05% Tween 20 and blocked with casein blocking buffer (37528 Thermo Fisher Scientific) containing 0.05% Tween 20. Serum samples were serially diluted from 1:100 to 1:1,968,300 in 3-fold steps and incubated for 1 hour at room temperature. Bound antibodies were detected with biotin-conjugated goat anti-mouse IgG, IgG1, IgG2a, or IgG2c antibodies (1030-08, 1070-08, 1080-08, or 1079-08, SouthernBiotech), followed by streptavidin–HRP (DY998, R&D Systems). Plates were developed with TMB substrate (34029, Thermo Scientific), and absorbance was measured at 450 nm using a Tecan Infinite M1000 Pro plate reader. IgG2c was measured in C57BL/6 mice, and IgG2a was measured in BALB/c mice. Results are reported as area under the curve calculated using GraphPad Prism.

### SARS-CoV-2 Spike and RBD ELISAs

SARS-CoV-2–specific antibody responses were measured by ELISA according to established protocols with minor modifications (*51*),(*49*),(*52*). High-binding 96-well MaxiSorp plates were coated overnight at 4°C with recombinant SARS-CoV-2 Spike or receptor-binding domain proteins corresponding to the WA1 or XBB.1.5 strains. After coating, wells were blocked for 1 hour at room temperature with PBS containing Tween-20 and milk powder. Serum samples were diluted in assay buffer, added to the plates, and incubated for 2 hours at room temperature. Plates were then washed with PBS-T. Bound antibodies were detected using HRP-conjugated anti-mouse IgG, IgG1, or IgG2c antibodies. Plates were developed with TMB substrate, and the reaction was stopped with sulfuric acid. Absorbance was measured at 450 nm with background correction at 570 nm using a SpectraMax i3 microplate reader. Results are reported as area under the curve calculated using GraphPad Prism.

### Influenza microneutralization assay with NP ELISA readout

Functional neutralizing activity of mouse sera was measured using a live influenza microneutralization assay with influenza A/California/07/2009 (H1N1) virus (VR-1894, ATCC), as previously described (*53*). Virus input was standardized to 100 TCID₅₀ per 50 µL, as determined in MDCK.2 cells (CRL2936, ATCC). Serial serum dilutions from 1:30 to 1:15,360 were prepared in 2-fold steps and preincubated with virus for 1 hour at room temperature. Serum–virus mixtures were then added to confluent MDCK.2 cell monolayers. After 1 hour, the inoculum was removed and replaced with infection medium containing the corresponding serum dilution. Cells were incubated for 72 hours at 37°C, after which culture supernatants were collected for quantification of influenza nucleoprotein. Influenza A NP levels in culture supernatants were measured using a commercial NP ELISA Pair Set (SEK004, Sino Biological). Clear 96-well Nunc MaxiSorp plates (439454, Thermo Scientific) were coated with rabbit anti-NP capture antibody at 0.5 µg/mL and detected with HRP-conjugated mouse anti-NP antibody. Recombinant NP standards ranging from 4,000 to 62.5 pg/mL were used to generate standard curves. Absorbance was measured at 450 nm using a Tecan Infinite M1000 Pro plate reader. Neutralization titers were defined as the reciprocal of the highest serum dilution that completely inhibited viral replication, as indicated by undetectable NP signal. A titer of ≥1:40 was used as a benchmark threshold (*54*).

### SARS-CoV-2 pseudovirus neutralization assay

Vero E6-ACE2/TMPRSS2 cells (2 × 10⁴ per well) were seeded in 96-well plates 24 hours prior to infection. Serum samples were heat-inactivated for 30 min at 56 °C and initially diluted 1:50 against the USA-WA1/2020 pseudovirus, followed by up to eight twofold serial dilutions. Diluted sera were incubated with pseudovirus preparations for 1 h at 37 °C with 5% CO₂. After incubation, the culture medium was aspirated and replaced with 50 µl of the serum–virus mixture. At 24 hours post-infection, luciferase reporter activity was measured using the Renilla-Glo Luciferase Assay System (Promega). Cells were lysed in 50 µl of lysis buffer, and 15 µl of lysate was mixed with an equal volume of luciferase substrate reagent. Luminescence was recorded using SpectraMax i3 microplate reader (Molecular Devices). Neutralization curves were analyzed in GraphPad Prism 10 (GraphPad Software) using nonlinear regression to determine the median inhibitory concentration (IC₅₀). Neutralizing antibody titers (NT₅₀) were defined as the reciprocal value of the IC₅₀.

### SARS-CoV-2 lung viral load quantification

To determine viral load following challenge experiments, lungs were collected in 2-ml screw-cap tubes (MP Biomedicals) containing DMEM (Gibco) supplemented with 2% FBS (Corning) and homogenized with Lysing Matrix D (MP Biomedicals) using a FastPrep-24™ Classic Bead Beating Grinder and Lysis System (MP Biomedicals). Samples were centrifuged, and the resulting supernatants were serially diluted seven times in 1:10 dilution ratio, starting with 200 µl of undiluted supernatant (20 µl sample in 180 µl medium, repeated seven times). A 25-µl aliquot from each dilution was added to six replicate 96-well plates containing Vero E6-ACE2/TMPRSS2 cells at 80–90% confluency in a monolayer. Cells had been seeded 48 hours earlier at a density of 2.4 × 10⁴ cells per well in 50 µl of DMEM (Gibco) supplemented with 2% FBS (Corning) and maintained at 37 °C with 5% CO₂. For the TCID₅₀ assay, each sample was distributed across six replicate plates (12 samples per plate). On the day of inoculation, an additional 50 µl of fresh DMEM (Gibco) supplemented with 2% FBS (Corning) was added to each well. After 5 days of incubation, cells were examined microscopically for signs of infection, including cytopathic effect (CPE). Viral titers (TCID₅₀) were calculated using the Spearman–Kärber method.

### Flow cytometry

Spleens and mediastinal lymph nodes were harvested, and single-cell suspensions were generated using a syringe plunger and a 70-µm cell strainer. Red blood cells were lysed with RBC Lysis Buffer. Cells were homogenized and resuspended in PBS containing 2% FBS and 0.5 mM EDTA for counting and staining.

For B-cell staining, cell suspensions were incubated with Fc block and LIVE/DEAD Fixable Near-IR Viability Kit, followed by antibodies against CD3, B220, CD19, CD38, CD45.2, CD95, CD138, GL7, IgD, and IgM.

For T-cell staining, cell suspensions were incubated with Fc block, LIVE/DEAD Fixable Near-IR Viability Kit, anti-CXCR5, and SARS-CoV-2 Spike-specific MHC tetramers provided by the NIH Tetramer Core Facility. A secondary incubation was then performed with antibodies against B220, CD3, CD4, CD8α, CD44, CD45.2, CXCR3, PD-1, and CD162.

Cells were fixed with 4% paraformaldehyde and resuspended in PBS containing 2% FBS and 0.5 mM EDTA. All samples were passed through a 70-µm cell strainer before acquisition on a BD FACSymphony A5 flow cytometer. Data were analyzed using FlowJo software version 10.100.0.

The complete antibody panel, including clones, vendors, and catalog numbers, is provided in **Supplementary Table S3**.

### RNA isolation, sequencing, and transcriptomic analysis

Injected muscle, liver, and lung tissues were collected from 9-week-old female C57BL/6 mice 6 h and 24 h after intramuscular administration of LNP formulations containing FL711, mRNA-Spike, or their combination, as described in the text and figure legends. Untreated tissue-matched samples from age- and sex-matched control animals were used as controls. For each condition, RNA from three mice was pooled before sequencing. RNA-seq libraries were prepared using a KAPA whole-transcriptome ribo-depletion workflow and sequenced on an Illumina NovaSeq platform in paired-end 100-bp mode. The transcriptomic analysis included LNP(FL711) alone, LNP(FL711) plus LNP(mRNA-Spike), LNP(mRNA-Spike) alone, and untreated controls from muscle, liver, and lung. Time points were analyzed separately.

Paired-end reads were quality-checked with FastQC (*55*) and processed with fastp using paired-end adapter detection, poly-G trimming, and a minimum retained read length of 30 nt (*56*). Trimmed reads were aligned to the mouse GRCm39 genome using STAR (*57*) in two-pass mode with GENCODE (*58*) vM38 annotation. The STAR index was generated from the GRCm39 primary assembly and GENCODE vM38 GTF annotation using sjdbOverhang 99. Gene-level counts were generated with featureCounts on exon features grouped by gene_id, using paired-end counting, proper-pair filtering, chimeric-fragment exclusion, MAPQ ≥10, and sample-specific strandedness inferred from aligned reads (*59*). Per-sample featureCounts outputs were combined into a gene-by-sample count matrix for downstream analysis.

Gene counts were filtered to remove genes with fewer than 10 total counts across retained samples and normalized with DESeq2 using a design of ∼1. Size factors were estimated with the poscounts method, and blind variance-stabilizing transformation was applied (*60*). Because RNA was pooled before sequencing, replicate-based per-gene differential-expression testing was not performed.

For IFN-I and NF-κB response estimation, per-gene ΔVST values were calculated by subtracting the tissue-matched untreated control from each treated sample. Mean ΔVST values were then calculated across predefined IFN-I-responsive or NF-κB-responsive gene sets for each tissue, formulation, and time point. The gene lists are provided in **Supplementary Figure S5**.

For pathway enrichment analysis, descriptive log2 fold changes were calculated from DESeq2-normalized counts using a pseudocount of 0.5. Treatment groups were compared with tissue-matched untreated controls. Genes were ranked by descriptive log2 fold change and analyzed by preranked GSEA (*61*) using fgsea (*62*). Pathway enrichment was reported as normalized enrichment score, with FDR-adjusted fgsea p values used to mark enriched pathways. Pathway gene sets were obtained from MSigDB (*63*) using msigdbr for Mus musculus; selected KEGG and Reactome inflammatory and innate-immunity pathways were used for focused pathway analysis.

Downstream analysis used R and Python. R packages included DESeq2, fgsea, msigdbr, AnnotationDbi, and org.Mm.eg.db; Python was used for integration of processed outputs and figure generation. The analysis environment included R 4.4 and Python 3.11.

### Statistical analysis

Quantitative experimental data were analyzed using GraphPad Prism software unless otherwise indicated. Group comparisons were performed using two-tailed Mann–Whitney U-tests unless otherwise specified in the figure legends.

Influenza antibody and SARS-CoV-2 ELISA responses are reported as area under the curve. Neutralization titers are reported as reciprocal endpoint titers for the influenza microneutralization assay or as NT₅₀ values for the SARS-CoV-2 pseudovirus neutralization assay.

For RNA-seq analyses, pooled RNA samples were used; therefore, replicate-based per-gene differential-expression testing was not performed. Transcriptomic data were analyzed descriptively using normalized expression values, ΔVST-based response scores, and preranked GSEA, as described above.

Data are shown as individual values with medians or as otherwise indicated in the figure legends. P values less than 0.05 were considered statistically significant.

## Authors’ contributions

ELA, IIG, and AVG conceptualized the approach. BI, AVV, ELA, IIG, and AVG designed the study. AVV, OMG, IIG, BI, KA, NMA, and ASG performed the experiments. IM conducted the bioinformatic analysis. AVV, BI, ELA, IIG, and AVG wrote the manuscript. All authors reviewed and edited the final manuscript.

## Acknowledgements

We thank Akiko Iwazaki for valuable discussions, Anton Komar for help with mRNA sequence optimization, Ivan Bespalov and Yanina Kozachok for technical assistance. This work was supported by a grant from Bill and Melinda Gates Foundation to Flag Bio, Inc. with a subcontract to BI, and a contract from NIAID to Flag Bio, Inc.

