## Supplemental Material for "mRNA-Encoded TLR5 Agonist as an Immune Adjuvant"

### Supplementary Materials

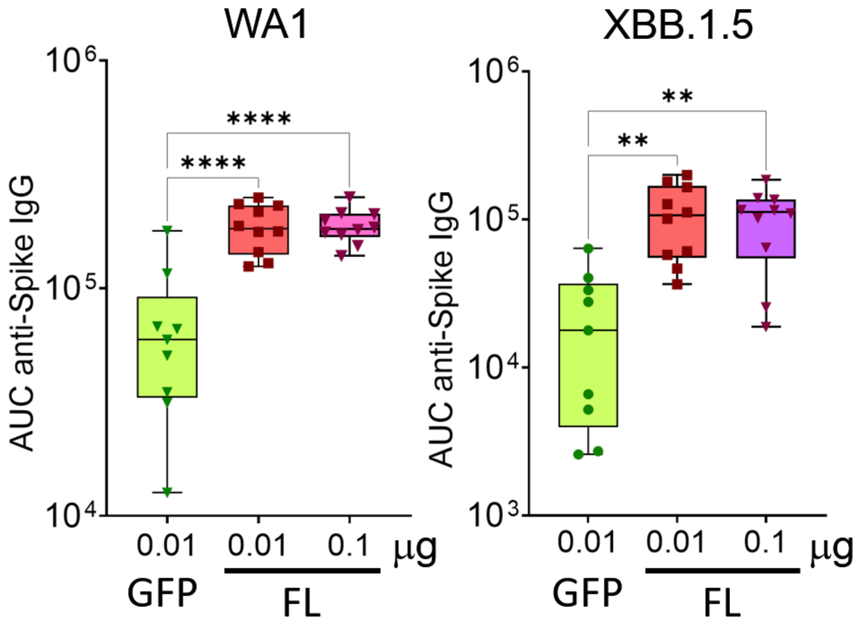


**Supplementary Figure S1. Similar adjuvating effects of 0.01 µg and 0.1 µg of LNP(FL711) when added to SARS-CoV-2 LNP(mRNA-Spike) vaccine.** As in Fig 2, K18-hACE2 mice were intramuscularly (IM) immunized (primed) with 0.3 µg of mRNA-LNPs encoding SARS-CoV-2 Spike protein (Hexapro, WA1) combined with either 0.01 µg of mRNA-LNPs encoding green fluorescent protein (GFP), 0.01 µg, or 0.1 µg of LNP( FL711). Mice received a booster immunization 21 days later with the same respective formulations. Serum was collected on day 56 post-prime (35 post-boost) to assess binding antibody responses against SARS-CoV-2 (WA1 or XBB.1.5) full Spike IgG. Boxes indicate median and interquartile range; whiskers show the full range; each point represents an individual mouse. Statistical significance was determined using the Mann–Whitney test. ns, not significant; $**P < 0.01$; $****P < 0.0001$.

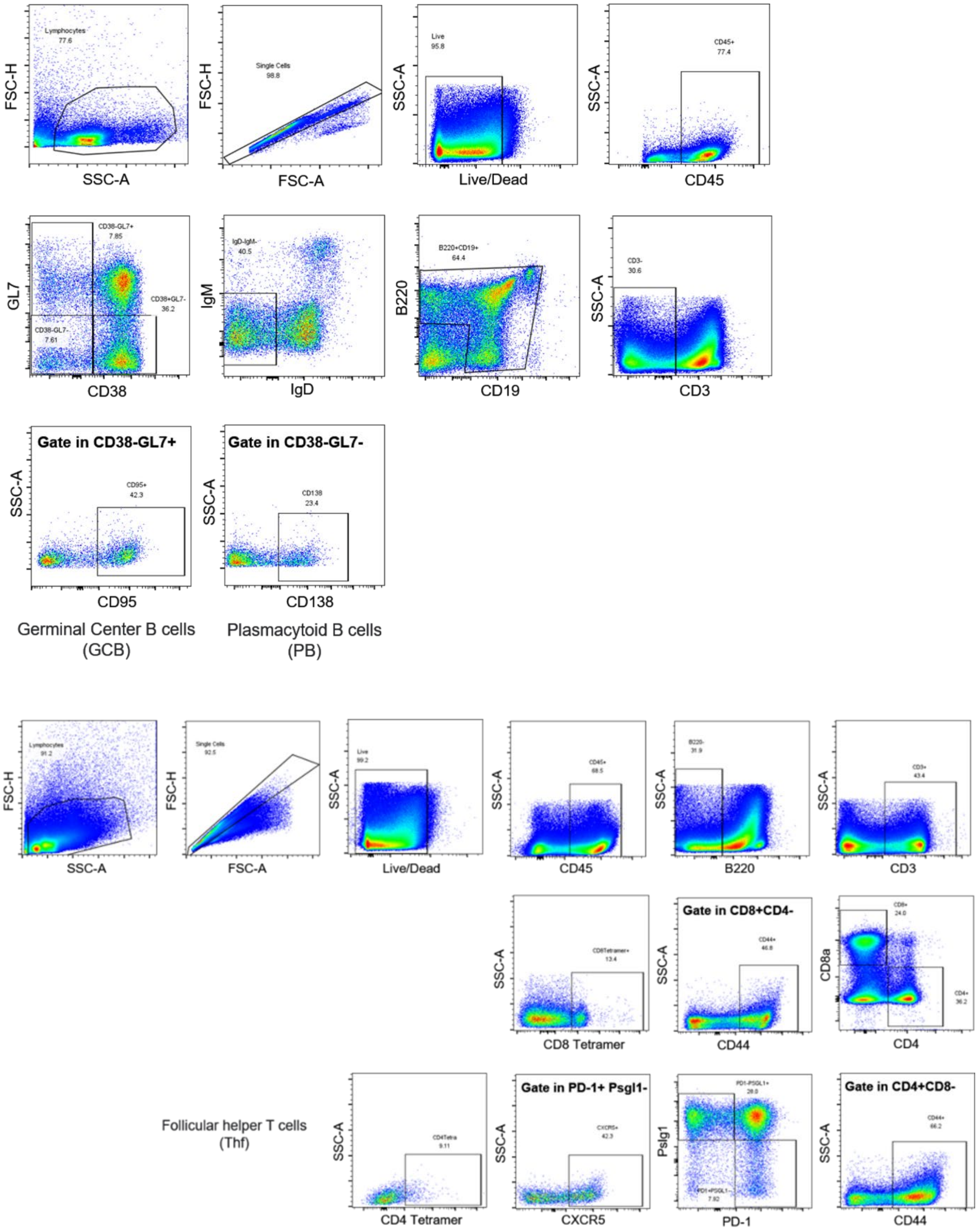


**Supplementary Figure S2. Gating strategies related to Fig. 2 J-M.** (A) Gating strategies to identify germinal center B cells and plasmablasts. (B) Gating strategies to identify antigen specific $CD4^{+}$ T cells, including $T_{FH}$ subsets.

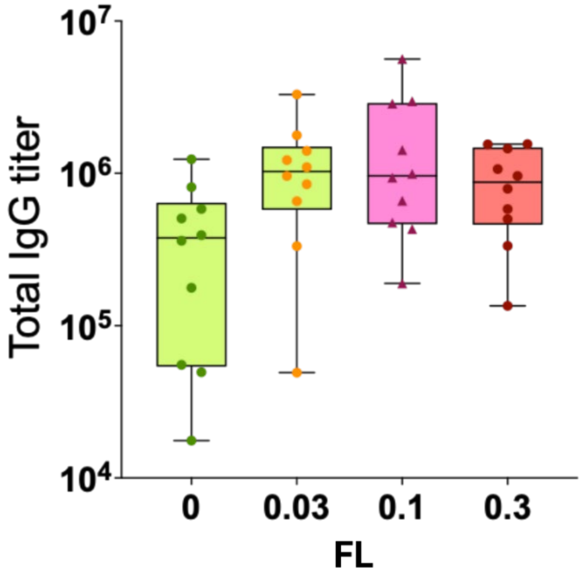


**Supplementary Figure S3. Dose selection of FL711 as an adjuvant for an influenza mRNA vaccine.** Total anti-HA IgG titers were measured in serum samples from BALB/c mice immunized with 0.5 µg per mouse of mRNA-LNPs encoding full-length hemagglutinin (HA) of influenza A/California/07/2009 (H1N1), administered alone or in combination with different doses of mRNA-LNPs encoding FL711. Individual symbols represent individual mice. Horizontal bars indicate the median, and error bars show the range. Although anti-HA IgG titers showed an increasing trend across FL711 dose groups, the overall group comparison by Kruskal–Wallis test did not reach statistical significance. The 0.3 µg dose of LNP(FL711) was selected for subsequent experiments because it produced the clearest enhancement of anti-HA IgG titers and was the only tested dose that significantly improved functional neutralizing antibody responses.

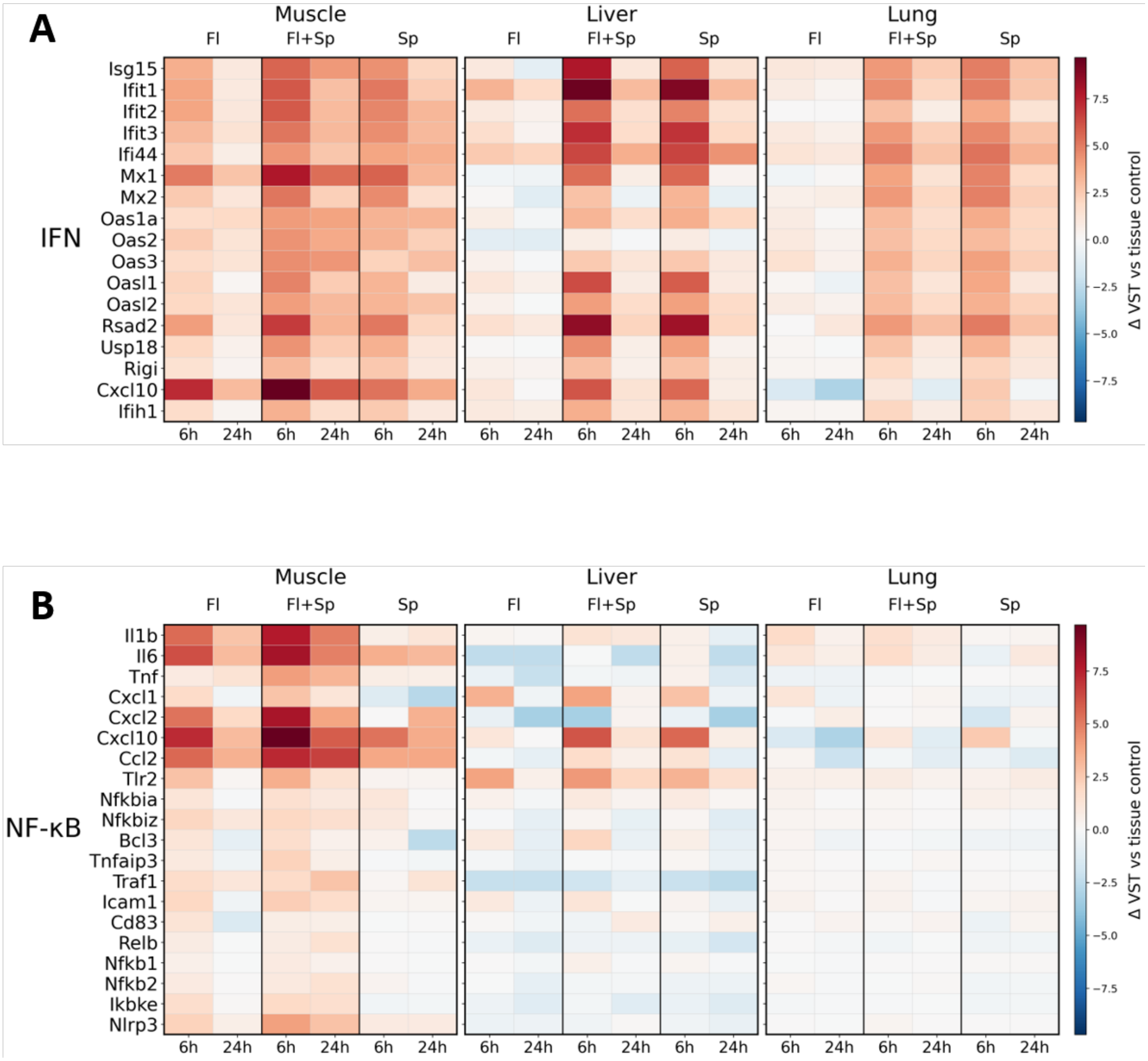


**Supplementary Figure S4.** Changes in expression of IFN-I-responsive **(A)** and NF-kB-responsive **(B)** gene sets in the indicated organs of mice following i.m. injection of three LNP(mRNA) vaccine preparations see legend to Figure 5 for details), including FL711 (Fl), FL711 + Spike (Fl+Sp) and Spike protein (Sp) of SARS-CoV-2 virus at two time points following injection. Heatmap values represent per-gene ΔVST expression relative to tissue-matched untreated controls, with common color scaling used within each panel.

**Supplementary Table S1.** Representative type I interferon–responsive genes selected for analysis (ISG panel), encompassing antiviral effectors, pattern-recognition receptors, chemokines, and feedback regulators that collectively illustrate the breadth of canonical IFN-I signaling.

| Gene | Full name | Canonical IFN-I responsiveness / role |
|---|---|---|
| ***Isg15*** | ISG15 ubiquitin-like modifier | Strongly induced ISG; conjugates to target proteins (ISGylation) and modulates antiviral and immune signaling. |
| ***Ifit1*** | Interferon-induced protein with tetratricopeptide repeats 1 | Robust ISG; binds cap0 viral RNAs and restricts translation of non–self mRNAs. |
| ***Ifit2*** | Interferon-induced protein with tetratricopeptide repeats 2 | IFN-inducible; contributes to antiviral defense and can promote apoptosis in infected cells. |
| ***Ifit3*** | Interferon-induced protein with tetratricopeptide repeats 3 | Strong ISG; forms complexes with other IFITs and stabilizes antiviral IFIT1/2 functions. |
| ***Ifi44*** | Interferon-induced protein 44 | Classical ISG; marker of IFN-I exposure, implicated in antiviral and antiproliferative responses. |
| ***Mx1*** | MX dynamin-like GTPase 1 | Potent IFN-inducible antiviral effector; blocks replication of many RNA viruses. |
| ***Mx2*** | MX dynamin-like GTPase 2 | IFN-inducible; restricts a subset of viruses (e.g., HIV-1, some DNA viruses) in a cell-type–dependent manner. |
| ***Oas1a*** | 2′-5′-oligoadenylate synthetase 1A | Strong ISG; synthesizes 2–5A to activate RNase L and degrade viral RNA. |
| ***Oas2*** | 2′-5′-oligoadenylate synthetase 2 | IFN-inducible; part of the OAS–RNase L antiviral RNA degradation pathway. |
| ***Oas3*** | 2′-5′-oligoadenylate synthetase 3 | Highly IFN-responsive; key activator of RNase L, often more potent than OAS1/2. |
| ***Oasl1*** | 2′-5′-oligoadenylate synthetase-like 1 | ISG; modulates IFN signaling, reported to negatively regulate IRF7 translation in mice. |
| ***Oasl2*** | 2′-5′-oligoadenylate synthetase-like 2 | IFN-inducible; antiviral effector with RNase L–independent antiviral activities. |
| ***Rsad2*** | Radical S-adenosyl methionine domain–containing 2 (Viperin) | Strong ISG; broadly antiviral via perturbation of lipid metabolism and viral budding/replication. |
| ***Usp18*** | Ubiquitin specific peptidase 18 | ISG; de-ISGylating enzyme and key negative regulator of IFN-I signaling (desensitization). |
| ***Rigi (Ddx58)*** | DExD/H-box helicase 58 (RIG-I) | IFN-inducible PRR; senses short 5′-triphosphate viral RNAs and triggers IFN production. |
| ***Cxcl10*** | C-X-C motif chemokine ligand 10 (IP-10) | Strongly IFN-inducible chemokine; recruits $CXCR3^+$ T cells and NK cells to inflamed tissues. |
| ***Ifih1*** | Interferon induced with helicase C domain 1 (MDA5) | IFN-inducible PRR; senses long dsRNA and promotes type I IFN production. |

**Supplementary Table S2.** Representative NF-κB–responsive genes encompassing cytokines, chemokines, receptors, transcriptional regulators, and feedback inhibitors that collectively illustrate the breadth of canonical inflammatory NF-κB signaling.

| Gene | Full name | Canonical NF-κB responsiveness / role |
|---|---|---|
| ***Il1b*** | Interleukin 1 beta | Strong NF-κB target; pro-inflammatory cytokine driving fever and leukocyte activation. |
| ***Il6*** | Interleukin 6 | Potent NF-κB-inducible cytokine; mediates acute-phase and systemic inflammatory responses. |
| ***Tnf*** | Tumor necrosis factor α | Core NF-κB-dependent cytokine; central to inflammation and apoptosis signaling. |
| ***Cxcl1*** | C-X-C motif chemokine ligand 1 | NF-κB-inducible neutrophil chemoattractant; early inflammatory marker. |
| ***Cxcl2*** | C-X-C motif chemokine ligand 2 | Similar to CXCL1; NF-κB-driven neutrophil recruitment. |
| ***Cxcl10*** | C-X-C motif chemokine ligand 10 | IFN-γ and NF-κB-responsive chemokine; attracts activated T cells. |
| ***Ccl2*** | C-C motif chemokine ligand 2 | NF-κB-inducible monocyte chemoattractant; hallmark of tissue inflammation. |
| ***Tlr2*** | Toll-like receptor 2 | NF-κB-inducible PRR; amplifies innate immune signaling. |
| ***Nfkbia*** | NF-κB inhibitor α (IκBα) | Immediate-early NF-κB target; negative feedback regulator of pathway activity. |
| ***Nfkbiz*** | NF-κB inhibitor ζ (IκBζ) | Inducible co-activator; modulates secondary NF-κB transcriptional waves. |
| ***Bcl3*** | B-cell lymphoma 3 | NF-κB-responsive nuclear IκB family member; fine-tunes transcriptional output. |
| ***Tnfaip3*** | TNF-α-induced protein 3 (A20) | Strong NF-κB target; ubiquitin-editing enzyme that terminates NF-κB signaling. |
| ***Traf1*** | TNF receptor-associated factor 1 | NF-κB-inducible adaptor; participates in TNFR and TLR signaling cascades. |
| ***Icam1*** | Intercellular adhesion molecule 1 | NF-κB-driven adhesion molecule; promotes leukocyte–endothelium interactions. |
| ***Cd83*** | CD83 molecule | NF-κB-inducible surface marker; upregulated during dendritic-cell activation. |
| ***Relb*** | RELB proto-oncogene, NF-κB subunit | NF-κB-responsive transcription factor; part of the non-canonical NF-κB pathway. |
| ***Nfkb1*** | NF-κB subunit p105/p50 | Transcriptionally regulated by NF-κB; forms canonical dimers. |
| ***Nfkb2*** | NF-κB subunit p100/p52 | Inducible component of non-canonical NF-κB signaling. |
| ***Ikbke*** | IκB kinase ε | NF-κB-inducible kinase; integrates antiviral and inflammatory signaling. |
| ***Nlrp3*** | NLR family pyrin domain-containing 3 | NF-κB-primed inflammasome sensor; links transcriptional activation to IL-1β maturation. |

**Supplementary Table S3. Flow-cytometry antibodies, tetramers, and reagents used for immune profiling.** Spleens and mediastinal lymph nodes were processed into single-cell suspensions and stained with the antibodies, tetramers, viability dye, and reagents listed below. Samples were acquired on a BD FACSymphony™ A5 flow cytometer and analyzed using FlowJo software version 10.100.0.

| Panel | Target / Reagent | Clone / Specificity | Supplier | Catalog No. | Use / Notes |
|---|---|---|---|---|---|
| B-cell and T-cell panels | Fc block | 2.4G2 | BD Biosciences | 553141 | Fc receptor blocking |
| B-cell and T-cell panels | LIVE/DEAD™ Fixable Near-IR Viability Kit | — | Thermo Fisher | L34980 | Viability staining |
| B-cell panel | CD3 | 17A2 | BioLegend | 100222 | T-cell exclusion / lineage marker |
| B-cell panel | B220 | HIS24 | BD Biosciences | 748446 | B-cell marker |
| B-cell panel | CD19 | 6D5 | BD Biosciences | 563557 | B-cell marker |
| B-cell panel | CD38 | 90/CD38 | BioLegend | 102741 | B-cell differentiation marker |
| B-cell panel | CD45.2 | 104 | BioLegend | 109806 | Leukocyte marker |
| B-cell panel | CD95 | SA367H8 | BioLegend | 152612 | Germinal center B-cell marker |
| B-cell panel | CD138 | 281-2 | BioLegend | 142519 | Plasmablast/plasma cell marker |
| B-cell panel | GL7 | GL7 | BioLegend | 144619 | Germinal center B-cell marker |
| B-cell panel | IgD | 11-26c.2a | BioLegend | 405725 | Naïve/mature B-cell marker |
| B-cell panel | IgM | RMM-1 | BioLegend | 406529 | B-cell isotype marker |
| T-cell panel | CXCR5 | L138D7 | BioLegend | 145517 | Tfh-associated marker; first incubation |
| T-cell panel | SARS-CoV-2 Spike S539–546 I-A$^b$ tetramer | VTWFHAIHVSGTNGT | NIH Tetramer Core Facility | 67914 | Spike-specific CD4+ T-cell detection |
| T-cell panel | B220 | RA3-6B2 | BioLegend | 103236 | B-cell exclusion / lineage marker |
| T-cell panel | CD3 | 17A2 | BioLegend | 100222 | T-cell marker |
| T-cell panel | CD4 | GK1.5 | BD Biosciences | 612923 | CD4+ T-cell marker |
| T-cell panel | CD8α | 53-6.7 | BioLegend | 100759 | CD8+ T-cell marker |
| T-cell panel | CD44 | IM7 | BD Biosciences | 751414 | T-cell activation/memory marker |
| T-cell panel | CD45.2 | 104 | BioLegend | 109806 | Leukocyte marker |
| T-cell panel | CXCR3 | CXCR3-173 | BioLegend | 126523 | Th1/Tc1-associated chemokine receptor |
| T-cell panel | PD-1 | 29F.1A12 | BioLegend | 135216 | Activation / Tfh-associated marker |
| T-cell panel | CD162 | 2PH1 | BioLegend | 746749 | PSGL-1; T-cell differentiation/activation marker |
| T-cell panel | SARS-CoV-2 Spike S539–546 H2-K$^b$ tetramer | VNFNFNGL | NIH Tetramer Core Facility | 75355 | Spike-specific CD8+ T-cell detection |
| Sample preparation | RBC Lysis Buffer | — | BioLegend | — | Red blood cell lysis |
| Sample preparation | 70-µm cell strainer | — | Research Products International | — | Preparation and filtering of single-cell suspensions |
| Fixation | Paraformaldehyde | 4% | Electron Microscopy Sciences | — | Cell fixation |
| Acquisition | BD FACSymphony™ A5 | — | BD Biosciences | — | Flow-cytometry acquisition |
| Analysis | FlowJo software | Version 10.100.0 | BD Biosciences | — | Flow-cytometry data analysis |

Abbreviations: medLNs, mediastinal lymph nodes; Tfh, T follicular helper; PSGL-1, P-selectin glycoprotein ligand-1.
